# Rapid macropinocytic transfer of α-synuclein to lysosomes

**DOI:** 10.1101/2022.01.06.475207

**Authors:** Armin Bayati, Emily Banks, Chanshuai Han, Wen Luo, Wolfgang E. Reintsch, Cornelia E. Zorca, Irina Shlaifer, Esther Del Cid Pellitero, Benoit Vanderperre, Heidi M. McBride, Edward A. Fon, Thomas M. Durcan, Peter S. McPherson

## Abstract

The nervous system spread of alpha-synuclein fibrils is thought to cause Parkinson’s disease (PD) and other synucleinopathies, yet the mechanisms underlying internalization and cellular spread are enigmatic. Here we use confocal and super-resolution microscopy, subcellular fractionation and electron microscopy (EM) of immunogold labelled alpha-synuclein preformed fibrils (PFF) to demonstrate that this fibril form of alpha-synuclein undergoes rapid internalization and is targeted directly to lysosomes in as little as 2 minutes. Uptake of PFF is disrupted by macropinocytic inhibitors and circumvents classical endosomal pathways. Immunogold-labelled PFF are seen at the highly curved inward edge of membrane ruffles, in newly formed macropinosomes, in multivesicular bodies and in lysosomes. While most fibrils remain in lysosomes, a portion is transferred to neighboring naïve cells along with markers of exosomes. These data indicate that PFF use a unique internalization mechanism as a component of cell-to-cell propagation.

## INTRODUCTION

A classic hallmark of Parkinson’s disease (PD) is the formation of Lewy Bodies (LBs). First discovered in 1912 (Lewy, 1912), LBs are cytoplasmic inclusions composed of fragments of membranous organelles and filamentous proteins (Shahmoradian et al., 2019; Tanaka et al., 2004). LBs are also observed in PD-related disorders such as Lewy body dementia. Alpha-synuclein (α-syn), encoded by the *SNCA* gene, is a major component of LBs and is implicated in their formation (Conway et al., 1998; Spillantini et al., 1997). α-syn appears to function in membrane trafficking and membrane curvature (Fortin et al., 2004; Jao et al., 2004; Nakamura et al., 2008; Vargas et al., 2017), and increased knowledge of its protein structure and the conformation of altered variants has led to enhanced understanding of α-syn misfolding and aggregation in disease (Lee et al., 2002; Li et al., 2002; Sandal et al., 2008).

Since the early 2000s, the Braak hypothesis has stated that PD develops with the spread of α-syn through the brain (Braak et al., 2003). This model gained traction with the observation that proteinaceous inclusions spread from brain tissue into implanted embryonic stem cells (Li et al., 2008), a result subsequently confirmed in animal models (Desplats et al., 2009b; Li et al., 2010; Recasens and Dehay, 2014; Recasens et al., 2018). Propagating pathology is also observed following injection of the fibril form of α-syn into localized brain regions of mice (Betemps et al., 2014; Luk et al., 2012; Masuda-Suzukake et al., 2014). This tendency to spread, along with the ability of α-syn fibrils to disrupt the conformation of endogenous α-syn, has earned the protein a label as a prion-like (Masuda-Suzukake et al., 2013; Mougenot et al., 2012). However, questions remain regarding the cell biological itinerary of α-syn propagation, notably the mode of cellular entry (Fenyi et al., 2019; Gelpi et al., 2014; Nakamura et al., 2015; Uemura et al., 2018; Yan et al., 2018).

Different mutations in the *SNCA* locus have varying penetrance that may correlate to the propensity of α-syn to aggregate (Lazaro et al., 2016; Rutherford et al., 2014). α-syn concentration is also a factor in aggregation as increased protein levels enhance aggregation, be it mutant or wild-type protein (Fink, 2006; Manning-Bog et al., 2002; Uversky, 2007). Further, α-syn preformed fibrils (PFF) are more effective at spreading and seeding α-syn aggregates than monomeric and other fibrillar forms (Conway et al., 2000; Lam et al., 2016). PFF comprise a heterogeneous number of α-syn monomers with various structural conformations (Pieri et al., 2016). Early studies using PFF revealed their ability to seed and induce PD pathology in cultured cells (Luk et al., 2009); hence, developing a protocol for consistent production of PFF was a significant contribution (Volpicelli-Daley et al., 2014; Volpicelli-Daley et al., 2011).

A key question in PD research relates to the means by which α-syn fibrils enter cells (Bieri et al., 2018). Several studies have concluded that α-syn endocytosis is clathrin-dependent, based on the use of dynamin and clathrin inhibitors, dynasore and pitstop, respectively, and clathrin heavy chain (CHC) colocalization (Konno et al., 2012; Liu et al., 2007; Rodriguez et al., 2018). It is generally thought that PFFs are then trafficked to lysosomes via the endosomal system over the course of tens of minutes to multiple hours. We found approximately 40 studies examining α-syn fibril endocytosis with variable conclusions (**Table S1**). Some of the variability may arise from examining α-syn internalization at longer time courses and not immediately after its addition to cells. Moreover, dynamin inhibition is not synonymous with inhibition of clathrin-mediated uptake and the specificity of dynasore and pitstop as inhibitors of clathrin-mediated endocytosis (CME) has come under scrutiny as both have off-target effects (Gu et al., 2010; Oh et al., 1998; Park et al., 2013; Pelkmans et al., 2002; Preta et al., 2015). Apart from CME, it is suggested that α-syn enter cells via direct permeation of the plasma membrane (Danzer et al., 2007), via the formation of tunneling nanotubes that allow direct connections between cells (Dieriks et al., 2017), or through caveolae-dependent endocytosis (Madeira et al., 2011). Thus, there appears to be no consensus nor definitive evidence regarding the nature of α-syn endocytosis.

We thus sought to examine endocytosis of PFF immediately after their addition to cells. Surprisingly, PFF are internalized rapidly and appear in lysosomes within 2 min, bypassing conventional endosomal trafficking pathways. We confirmed this result in multiple cell lines, primary human cells, and neurons derived from induced pluripotent stem cells (iPSC). The internalization is not dependent on clathrin but instead uses macropinocytosis. We used gold-labeled PFF and EM and discovered PFF in membrane ruffles that form macropinosomes and in lysosomes. We also detect gold-labeled PFF on the outer edges of inward invaginating vesicles in endosomes and on the outside of vesicles within multivesicular bodies (MVB). While a portion of PFF remain in lysosomes for a long period, a smaller portion are transferred to naïve cells along with markers of MVB. Thus, our data unveils a unique form of macropinocytosis that mediates internalization of PFF and allows for endocytosis to be coupled to release.

## RESULTS

### PFF are rapidly endocytosed to lysosomes

We used fluorescently labeled PFF (Del Cid Pellitero et al., 2019; Maneca et al., 2019) (**Fig. S1A/B**) with live-cell imaging and a trypan blue exclusion assay (Karpowicz et al., 2017) that quenches extracellular PFF fluorescence (**Fig. S1D**) to examine PFF internalization at the earliest possible time points. PFF are rapidly internalized in U2OS cells and are targeted to lysosomes where they colocalize with Lysosomal Cytopainter within as little as 2 min (**Fig. 1A**). Similar results are seen when incubating cells with PFF at 4°C for 30 min and then transferring the cells to 37°C (**Fig. S1E**). We confirmed the live imaging findings in fixed samples of U2OS cells. We confirmed the live imaging findings in fixed samples of U2OS cells. PFF are significantly internalized within 2 min and continue to accumulate within cells for up to 60 min (**Fig. 1B** **& S1H**). Nearly all labeled PFF that enter cells are colocalized with LAMP1-TurboRFP by 2 min, and this stays stable with all PFF trafficking to lysosomes) for up to 60 min (**Fig. 1C**). As fixation makes cells permeable to trypan blue, for fixed cell experiments, we used trypsin to proteolyze off extracellular PFF (**Fig. S1F/G**). The rapid uptake of PFF and transport to lysosomes within 2 min was also observed in U87 (**Fig. S2A/D**) and U251 (**Fig. S2B/E**) glioblastoma cells. U2OS cells continue to accumulate PFF for up to 72 h (**Fig. 1D**).

**Figure 1.**
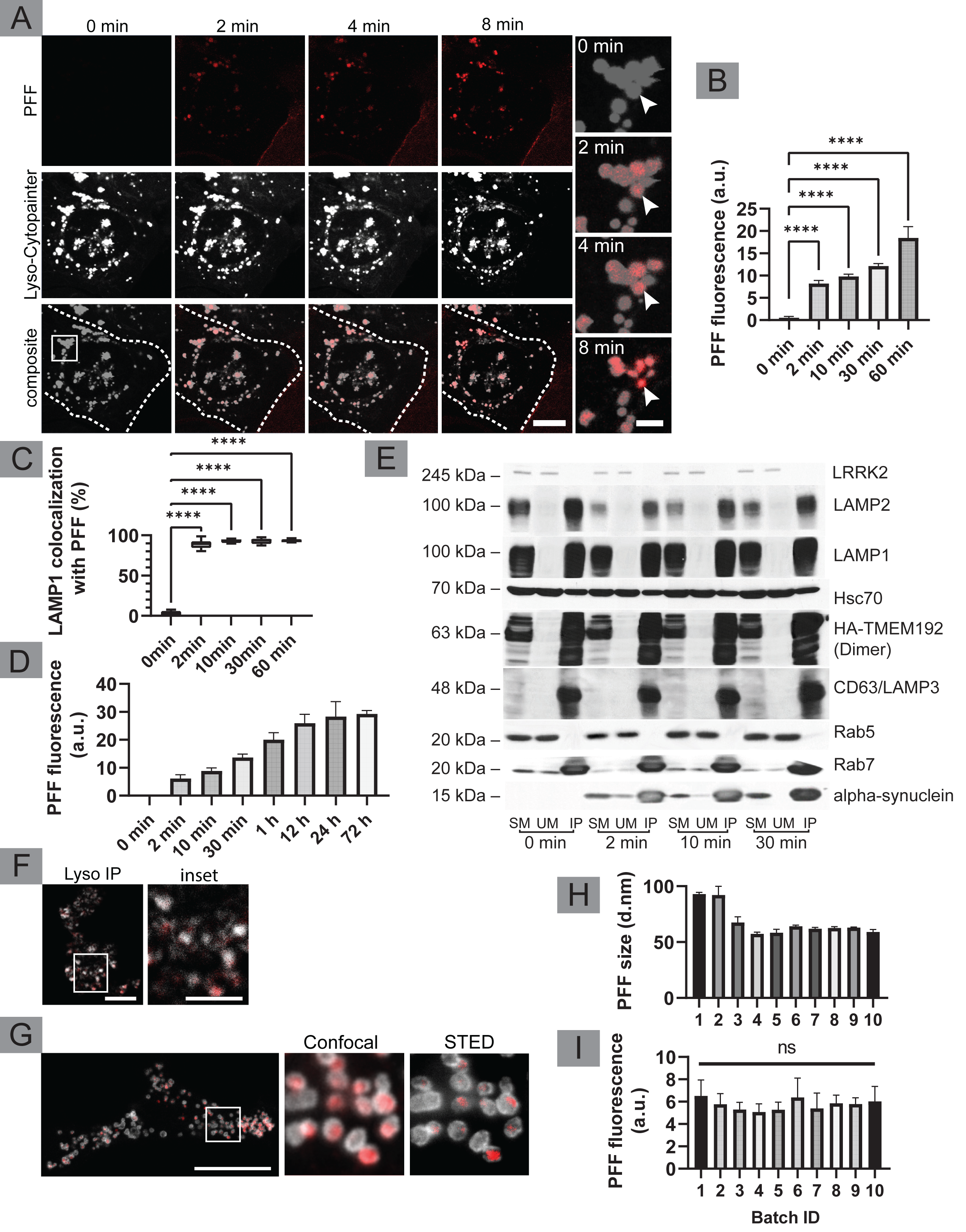
Rapid internalization and transport of PFF to lysosomes. (**A**) U2OS cells were stained with Lyso-Cytopainter and live-cell imaging was performed at 1 frame/sec. PFF tagged with Alexa488 at 2 µg/mL was added to cells while imaging. Colocalization of PFF (red) with lysosomes (white) can be seen as early as 2 min. Arrowheads point to lysosomes accumulating PFF. Scale bars = 10 µm for low mag images and 5 µm for insets. Inset location is shown at 0 min. (**B**) Quantification of PFF uptake in U2OS. Cells were plated on coverslips and transfected with LAMP1-TurboRFP. PFF were added to each coverslip at 2 µg/ml and cells were incubated for the indicated times following the addition of PFF. Cells were washed with trypsin and fixed. Colocalization of PFF with LAMP1 can be seen as early as 2 min; n = 12 for each condition (n = 60 total) for three independent experiments, mean ± SD. (**C**) Colocalization rate of LAMP1 with PFF from experiment detailed above in **A** and **B**, n = 12 for each condition (i.e., n = 60 total), from three independent experiments, mean ± min & max values. For **B** and **C**, one-way ANOVA and multiple comparisons Tukey’s test was conducted to compare means. p < 0.0001 denoted as ****. (**D**) Uptake of PFF in U2OS cells was quantified over the course of 72 h. Endocytosis of PFF plateaus at 24 h. n = 5 for each condition; mean ± SD. (**E**) Lysosomes were immunoprecipitated from U2OS cells stably expressing HA-TMEM192-RFP using HA-magnetic beads. Starting material (SM), unbound material (UB), and immunoprecipitated (IP) samples for each time point were collected and processed for Western blot with the indicated antibodies. (**F**) IP samples from the experiments as in **E** were imaged using confocal microscopes. HA-TMEM192-RFP construct allowed us to visualize the lysosomes (white), many of which contained PFF tagged with Alexa Fluor 488 (red). Scale bars = 5 µm for low mag and 2.5 µm for inset. (**G**) Confocal samples of U2OS cells described in **A** imaged using STED microscopy. Scale bar = 10 µm for low magnification, and 0.5 µm for insets. (**H**) Dynamic light scattering measurement of 10 different batches (3 different measurements using DLS for each batch) of conjugated PFF, were administered to U2OS cells and their uptake was measured in **I**. No significant difference was observed in the uptake of the different sizes of PFF. n = 9 for images analyzed corresponding to each batch (i.e. n = 90 total) from three independent experiments; mean ± SD, ns = not significant.

To confirm the rapid transport of PFF to lysosomes, we incubated U2OS cells stably expressing HA-TMEM192-RFP, a lysosomal protein, with PFF for various time periods, then lysed the cells and purified lysosomes using HA-magnetic beads (Abu-Remaileh et al., 2017). The enrichment of lysosomes was confirmed in the immunoprecipitated fractions with antibodies recognizing LAMP1 and 2 (**Fig. 1E**). Rab7 was also enriched in the lysosome fractions, whereas Rab5 and LRRK2 were not detected. CD63/LAMP3, a marker of lysosomes and MVB, was the most highly enriched marker. PFF were enriched in the lysosome fractions as early as 2 min, confirming their rapid transport to lysosomes (**Fig. 1E**). The enriched lysosomes were placed on coverslips and visualized through the fluorescently-labelled PFF (Alexa Fluor 488) and the RFP tag on lysosomes (**Fig. 1F**). PFF were detected in the lumen of the lysosomes, which was most readily seen using STED super-resolution microscopy (**Fig 1G**). Fluorescently conjugated PFF of various sizes (**Fig 1H**) were administered to U2OS cells revealing no significant difference in uptake in samples ranging from 50-100 nm (**Fig 1I**). Thus, PFF are rapidly endocytosed and transported to lysosomes in as little as 2 min, an unprecedented time frame for lysosomal targeting.

### Rapid transfer of PFF to lysosomes in nervous system cells including human dopaminergic neurons

Human cortical neurons derived from iPSCs (**Fig. S1I**) were incubated with fluorescently labeled PFF and colocalization with lysosomes was examined using a fixable form of Lysotracker. PFF were rapidly endocytosed in the neurons with lysosomal colocalization observed as early as 2 min (**Fig. S2C/F/G**), although lysosomes continued to accumulate PFF for up to 30 min. In both human dopaminergic neural progenitor cells (NPC) derived from iPSCs and in dopaminergic neurons derived from the NPC, PFF rapidly internalize and are detected at lysosomes within 2 min (**Fig. S1I, 2A, B, D, E**). Similar results are seen in human astrocytes (**Fig. S1I, 2C/F**). Thus, in multiple cell types, including those from the human nervous system, PFF are rapidly transported to lysosomes.

**Figure 2.**
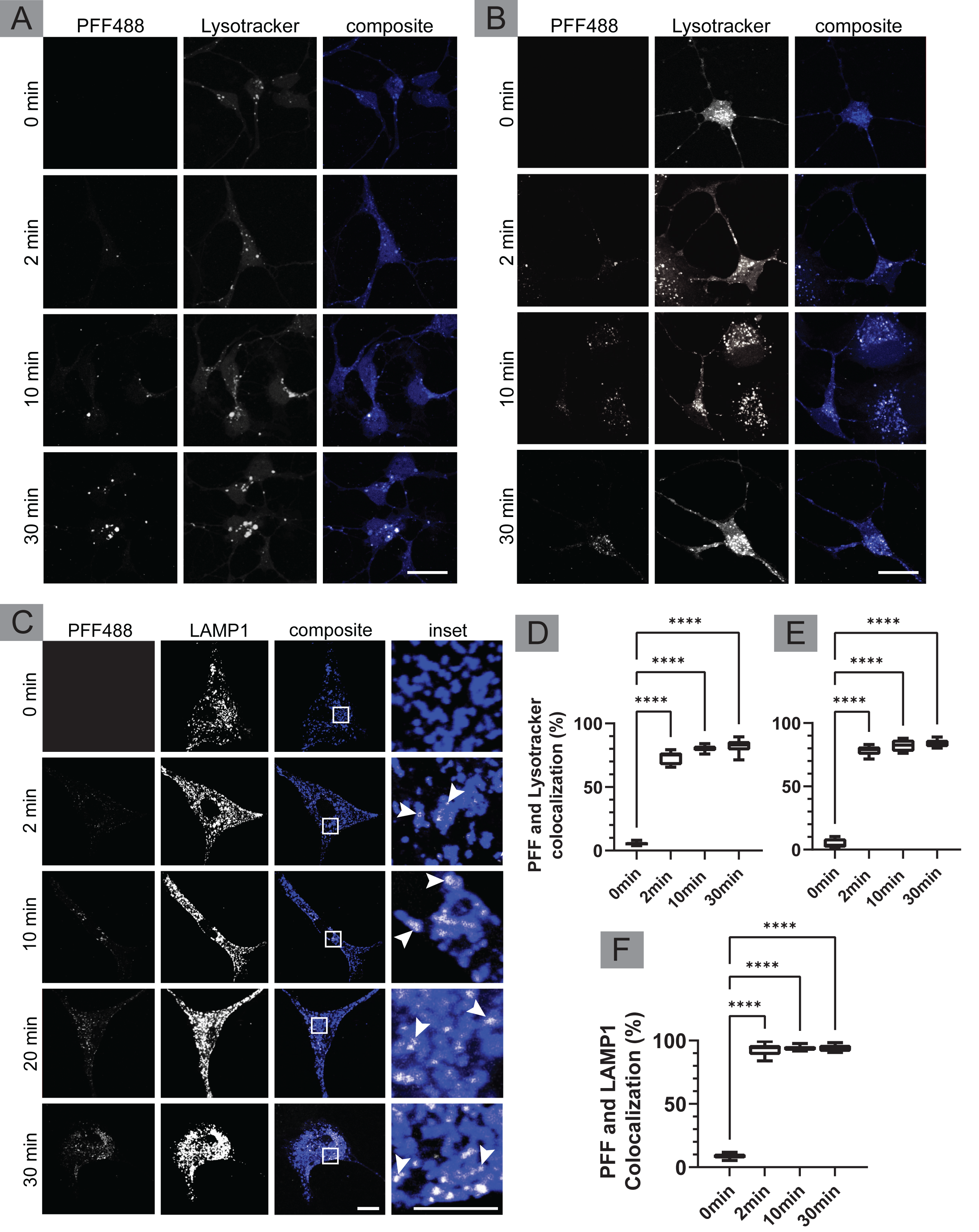
Rapid colocalization of PFF with lysosomes in human dopaminergic NPCs, dopaminergic neurons, and astrocytes. (**A**) Dopaminergic NPCs were incubated for 0, 2, 10, 30 min at 37°C following addition of Alex488-labelled PFF (white) at 2 µg/mL. The cells were then washed with trypsin, fixed, and stained with lysotracker (blue). Scale bar = 20 µm. (**B**) NPCs differentiated into dopaminergic neurons were processed as in **A**. Scale bar = 20 µm. (**C**) Human astrocytes were grown and mounted on coverslips, PFFs were added to each coverslip at 1 µg/ml, and cells were incubated for 0, 2, 10, and 30 min at 37°C following addition of Alex488-labelled PFF. Cells were washed with trypsin, fixed, permeabilized, and stained with LAMP1. Scale bar = 20 µm and 2.5 µm insets. (**D/E**) Colocalization of Lysotracker with PFF from experiments as in **A** and **B.** n = 6 for NPCs and n = 6 for neurons in each condition (n = 48 total for NPC and n = 48 for neurons), from three independent experiments. (**F**) Colocalization rate of LAMP1 with PFF from experiments as in **A.** n = 9 for each condition (i.e., n = 36 total), from three independent experiments. For all quantifications, one-way ANOVA was conducted followed by multiple comparisons Tukey’s test; mean ± min & max values; p < 0.0001 denoted as ****.

PFF were added to U343 glioblastoma cells for 24 h followed by a change to fresh media. PFF are trafficked to lysosomes and remain in the organelle, accumulating for up to 10 days (**Fig. S2J**). In contrast, internalized epidermal growth factor (EGF) dissipates from lysosomes (**Fig. S2J**). Internalized PFF remain in lysosomes in human dopaminergic neurons (**Fig. 3A**) and in human astrocytes (**Fig. 3B**), days after extracellular PFF has been removed from the cell media. The fluorescence intensity of PFF over 14 days was investigated in human astrocytes (**Fig 3C**). Quantification of the fluorescence showed a significant reduction in PFF fluorescence after day 1 (**Fig 3D**); however, a considerable amount of PFF remains in lysosomes at 14 days.

**Figure 3.**
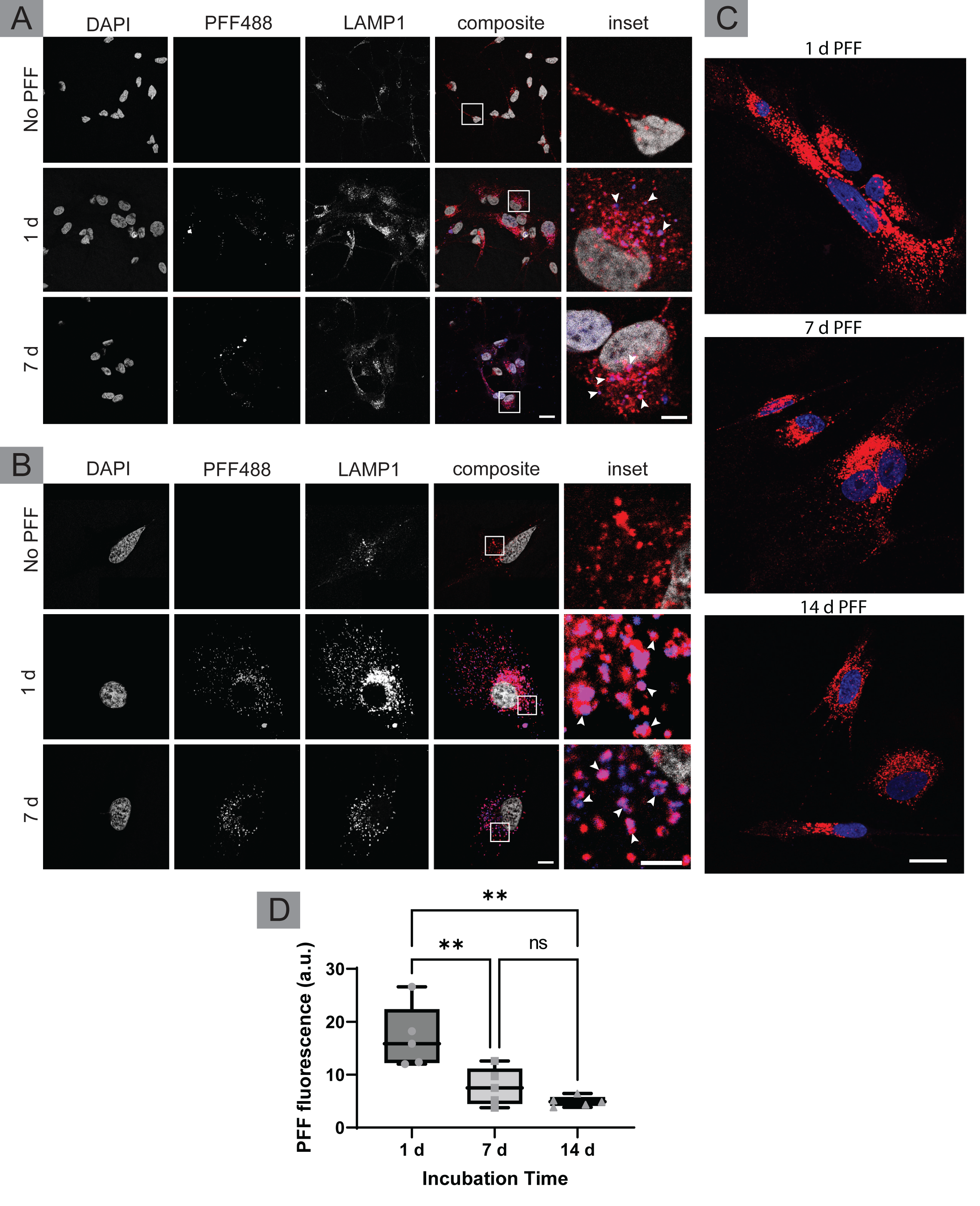
PFF remain in lysosomes for at least 7 days in dopaminergic neurons and astrocytes. (**A**) Dopaminergic neurons derived from human iPSCs mounted onto coverslips were given PFF488 at 2 µg/ml on ice for 1 h. Cells were removed from ice, given fresh media and were then incubated for 0 h, 1 day or 7 days at 37°C. Following incubation, cells were trypsin washed and fixed. Lysosomes were stained using LAMP1 (red) antibody. PFF (blue) remain localized to lysosomes 7 days following its addition to the cell. Scale bar = 20 µm for low magnification and 5 µm for insets. (**B**) Human astrocytes were mounted on coverslips and treated as described in **A**. PFF fluorescence builds in lysosomes over 7 days. Arrowheads point to LAMP1 and PFF colocalization. Scale bar = 20 µm for low magnification and 5 µm for insets. (**C**) Adult human astrocytes at passage 7 were plated on coverslips at 50% confluency and incubated with PFF for 24 h. The cells were then trypsin washed and given fresh media. Some cells were fixed while others were incubated for an additional 6 d or 13 d before fixation. Scale bar = 20 µm. (**D**) The intensity of PFF fluorescence was quantified for astrocytes as in **C**. n = 5 for each condition (n = 15 total) collected from five independent experiments. Data was then analyzed using one-way ANOVA, and Tukey’s test was conducted *post-hoc* to compare difference between means; mean ± min & max values; p < 0.001 depicted as **, p > 0.05 was set to be not significant = ns. Individual data points shown in gray.

### PFF are trafficked to lysosomes avoiding early/recycling endosomes

Considering the speed at which PFFs reach lysosomes it seems unlikely that PFF follow endosomal pathways to lysosomes (Lee et al., 2005; Lee et al., 2008; Lee et al., 2016) as endosomal maturation generally takes 10-15 min (Huotari and Helenius, 2011). Transferrin (Tf), a well-studied marker of early and recycling endosomes (Trischler et al., 1999) was added along with PFF and we observed no co-localization as 2 min and for as long as 30 min (**Fig. S3A/D**). Moreover, internalized PFF do not colocalize with EEA1, a marker of early endosomes (Mills et al., 1998), even at time points as early as 2 min (**Fig. S3B/E**). PFF colocalize with the late endosome/lysosome markers Rab9 and LAMP1 but show little co-localization with early and recycling Rabs, 4, 5 and 8 (**Fig.S3C, F/G**). Thus, PFF appear to reach lysosomes independently of the early and recycling endosomal systems.

### Lysosomal transfer of PFF does not depend on bulk endocytosis

We then moved to other markers of endocytosis to explore the identity of PFF internalization. Due to the rapid transport of PFF to lysosomes, we also sought to determine whether phagocytosis may be responsible for uptake of PFF. RAW 264.7 macrophage cells were activated with LPS (1 μg/ml) 24 h prior to the experiment. Cells were given fluorescent latex beads (FluoSpheres) with PFF. Little colocalization was observed between FluoSpheres and PFF (**Fig S4A**). We then used dextrans of different molecular weights were administered to U2OS cells alongside or prior to the addition of PFF (**Fig S4B**). We then moved to a macropinocytic marker, dextran (**Fig S4C**). Dextran with different molecular weights were used, due to the relationship between the size of dextran and its endocytic pathway (Li et al., 2015). The colocalization was only 25% when 10k dextran was added simultaneously with PFF (**Fig S4D**). Higher molecular weight dextran samples showed lower colocalization when added simultaneously with PFF. Thus PFF do not utilize bulk entry but seem to be internalized through a unique form of endocytosis.

### Endocytosis of PFF is clathrin-independent

The current consensus is that PFF enter cells via CME (Uemura et al., 2020). However, we are aware of no mechanism by which cargo that enters cells via CME can gain access to lysosomes in 2 min while bypassing early endosomes. To test if CME is involved in the endocytosis of PFF, we used an established genetic approach involving the knockdown of CHC with previously validated siRNA (Galvez et al., 2007; Kim et al., 2011). Immunoblot reveals effective CHC knockdown in U2OS cells (**Fig. 4A**). We then plated cells treated with control siRNA or CHC-targeting siRNA as a mosaic in the same well and incubated them with PFF. We observed no difference in internalization of PFF when comparing knockdown and control cells (**Fig. 4B/C**). Thus, it does not appear that CME is a major route for the internalization of PFF.

**Figure 4.**
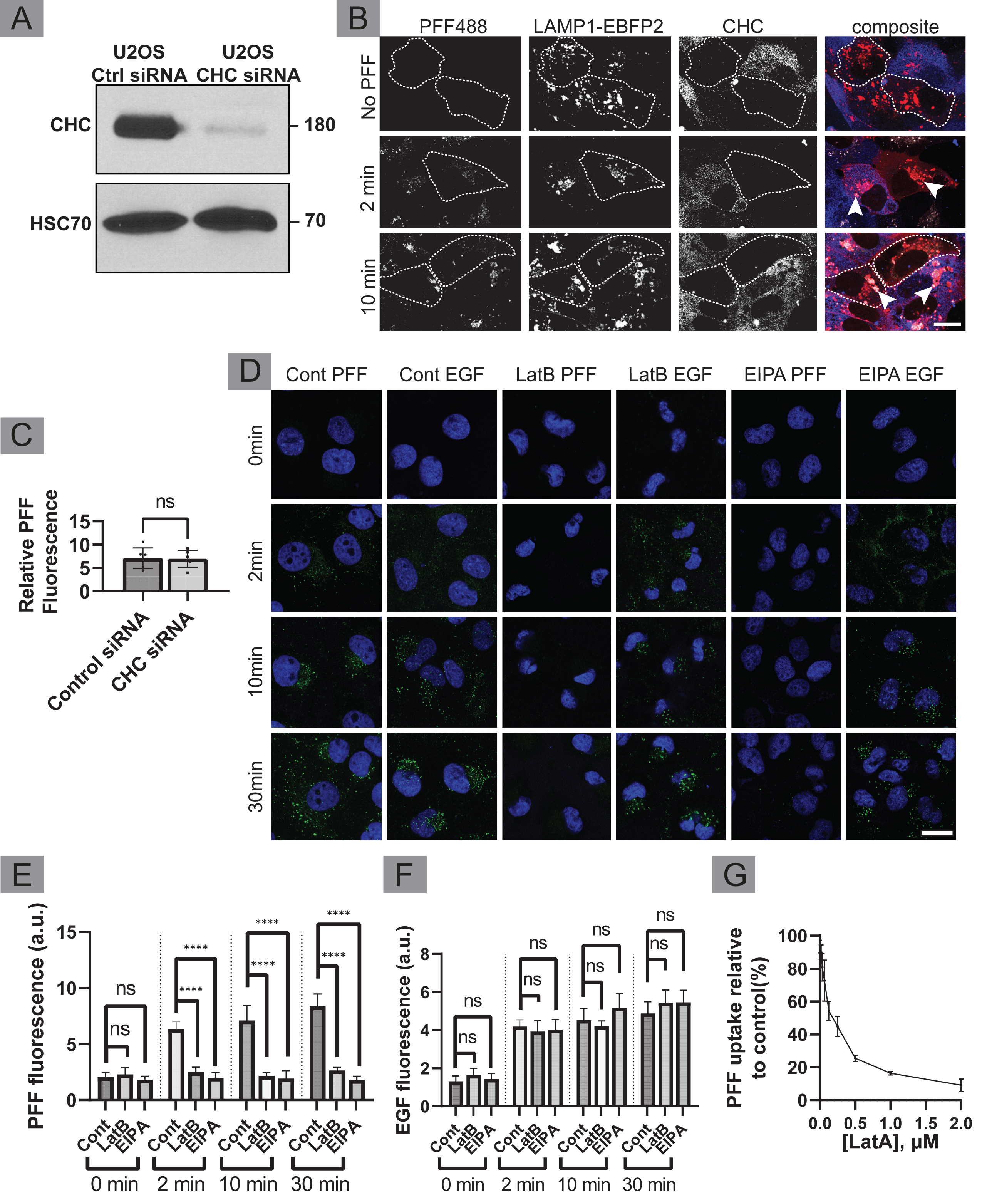
Endocytosis of PFF is unaffected by CHC KD but is decreased by macropinocytosis inhibitors. (**A**) U2OS cells previously transfected with EBFP2-LAMP1 were transfected with CHC siRNA or control siRNA. Cell lysates were immunoblotted with antibodies recognizing the indicated proteins (**B**) At 24 h, control and CHC siRNA-treated cells were re-plated as a mosaic onto coverslips. PFF were added to each coverslip at 2 µg/ml. Cells were incubated for 0, 2, 10 at 37°C following addition of PFF. Cells were washed with trypsin and fixed. Arrowheads show large LAMP1(red)- and PFF(white)-positive vesicles in both CHC (blue) positive and CHC negative cells, outlined by the CHC antibody. KD cells are outlined with dashed lines. Scale bar = 20 µm. (**C**) Internalization of PFF in CHC KD vs control cells at 2 min were quantified from experiments as in **B**, except that for quantification, KD cells were plated on separate coverslips than control. n = 6 for each condition (i.e., n = 12 total), from three independent experiments, mean ± SEM; unpaired t-test; p > 0.05 denoted as ns for not significant. (**D**) Astrocytes were grown and mounted on coverslips. Cell media was replaced with serum-free media containing 20 µM EIPA, 5 µM LatB or DMSO (vehicle control) for 30 min. PFF (green) was added to each coverslip at 2 µg/ml for 0, 2, 10, 30 min at 37°C. Cells were trypsin washed and fixed. Cell nuclei were stained with DRAQ7 (blue). Scale bar = 20 µm. (**E**) Quantification of PFF uptake from experiments as in **D**. At each timepoint, control was compared to LatB and EIPA using unpaired samples t-test. (**F**) Quantification of EGF uptake from experiments as in **D**. n=8 for each condition (n = 96 total for PFF and n = 96 for EGF), from three independent experiments, mean ± SD; p < 0.0001 denoted as ****, ns = not significant. (**G**). Human dopaminergic NPCs were treated with LatA and PFF for 24 h before fixation. y-axis: % intracellular PFF-Alexa fluorescence relative to DMSO (vehicle) only control (DMSO=100%). X-axis: Latrunculin A treatment concentration in µM; mean ± SEM

### Internalization of PFF occurs through macropinocytosis

Holmes et al. (2013) demonstrated that tau fibrils utilize macropinocytosis for cellular entry and that fibril α-syn colocalizes with tau during uptake, suggesting a potential role for macropinocytosis in the internalization of fibril α-syn. Consistently, Zeineddine (2015) (Zeineddine et al., 2015) found that α-syn fibrils induce membrane ruffling, an early step in forming macropinosomes. Although macropinocytic cargo generally follow the endosomal pathway (Mayor and Pagano, 2007), we sought to test if macropinocytosis is involved in the internalization of PFF. We first examined if EIPA, an established inhibitor of macropinocytosis (Commisso et al., 2014; Koivusalo et al., 2010; Nakase et al., 2015) influences the internalization of PFF. EIPA disrupts the Na^+^/H^+^ exchanger, decreasing cytosolic pH and inhibiting the activation of Cdc42 and Rac1, required for macropinocytosis (Koivusalo *et al*., 2010). At a concentration of 20 µM, EIPA inhibits uptake of PFF in human astrocytes (**Fig. 4D/E**). Latrunculin B (LatB), which inhibits macropinocytosis by disrupting actin polymerization (Williams and Kay, 2018), had a similar block on uptake of PFF when used at 5 µM (**Fig. 4D/E**). In contrast, neither drug influenced the uptake of EGF (**Fig. 4D/F**), which at the concentration used, enters cells via CME (Sigismund et al., 2005). In addition to astrocytes, we confirmed our findings in dopaminergic NPCs. Latrunculin A (LatA), which like LatB is a potent actin polymerization inhibitor (Fujiwara et al., 2018), demonstrated a dose-dependent inhibition of PFF uptake in dopaminergic NPCs with uptake reduced by 81% compared to control at 2 μM (**Fig. 4G**).

Consistent with a role for macropinocytosis in endocytosis of PFF, PFF induce the formation of actin-rich membrane ruffles, precursors of macropinosomes (Condon et al., 2018) (**Fig. S5A/B**). More specifically, PFF induce recruitment of Rac1 to actin-rich membranes on the cell surface, a characteristic of macropinocytosis (Grimmer et al., 2002) (**Fig. S5A**). The ability of PFF to induce membrane ruffling is also readily observed by EM (**Fig. S5C**). As a control, we examined the influence of EGF, a documented inducer of membrane ruffles, even though it does not use macropinocytosis itself for internalization (Bryant et al., 2007). We also examined Tf, which is not known to induce membrane ruffling or macropinocytosis. As expected, EGF causes the recruitment of Rac1 to the surface, where it colocalizes with F-actin, whereas no Rac1 recruitment is seen upon the addition of Tf (**Fig. S5D**). Thus, PFF appear to stimulate membrane ruffles and use macropinocytosis to gain direct access to lysosomes.

### Trafficking itinerary of PFF revealed by immunogold EM

To observe the trafficking itinerary of PFF directly we labeled the fibrils with gold (**Fig. S1C**) and followed their trafficking by EM. At 2-3 min following addition of PFF, astrocytes were fixed and processed for EM with uranyl acetate staining. Gold-labeled PFF appear under membrane ruffles and intracellular macropinosomes formed by ruffle closure (**Fig. 5Ai/ii**). The immunogold-labeled PFF remain in the lumen of the macropinosomes/endosomes that begin to demonstrate inward invagination of vesicles (**Fig. 5Aiii/iv**). At 3-5 min the PFF can be found in the lumen of intracellular membranes that gradually acquire electron density, indicating that they are likely lysosomes (**Fig. 5C**; **Fig. S6A**/**B**).

**Figure 5.**
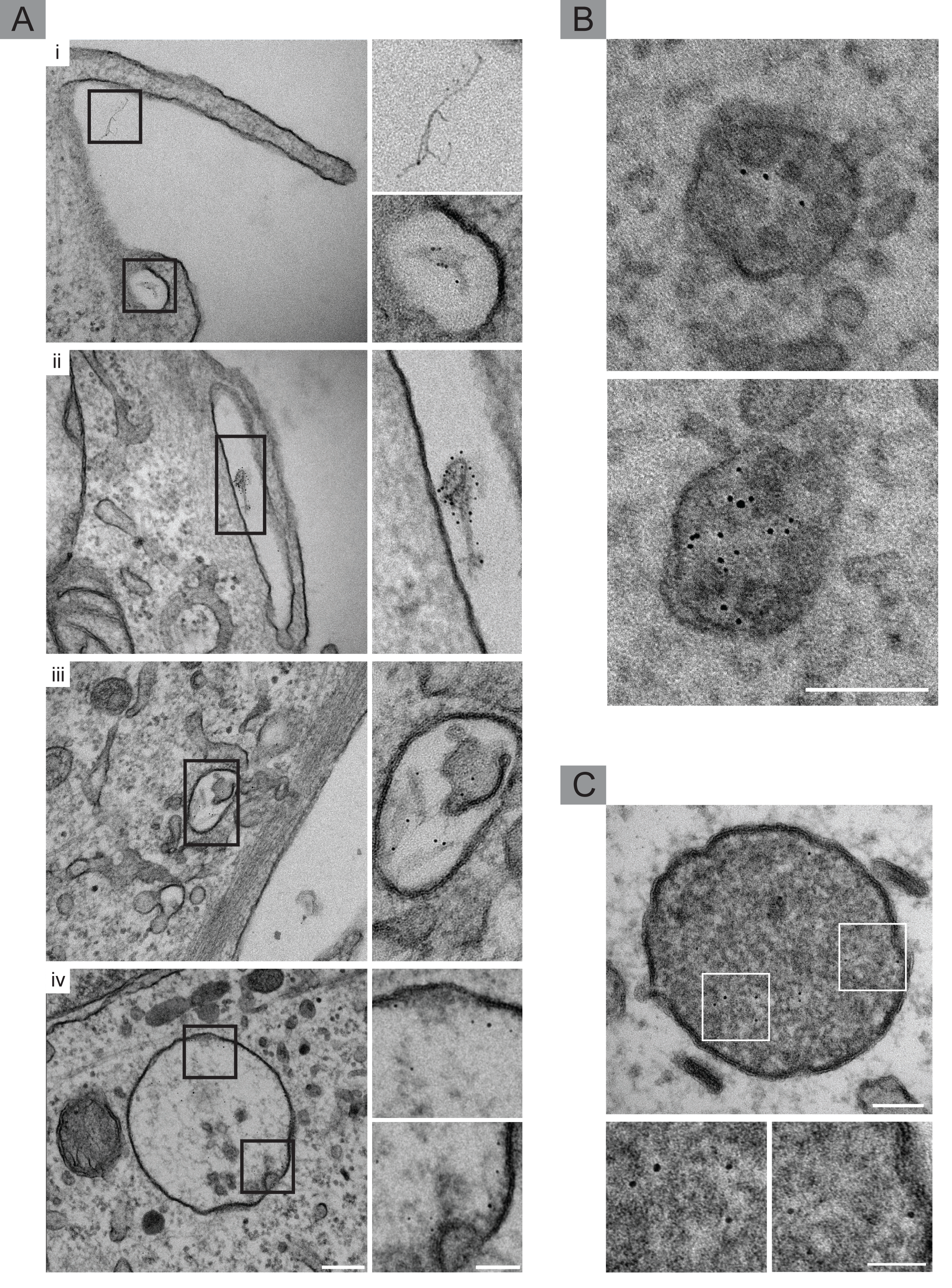
Trafficking itinerary of PFF revealed by nanogold-labelled PFF and EM. (**A**) Early events involved in the internalization of PFF were captured with EM. To allow direct detection, PFF were conjugated with 5 nm gold. Astrocytes were plated on permanox 8 well plates, administered PFF-gold for 2-3 min, fixed, and processed for EM. Uranyl acetate staining was conducted both pre-embedding and post-embedding (on the grid). (**i**) shows the membrane protrusion/ruffle forming around PFF-gold and the subsequent closure of the ruffle, with an enclosed macropinosome in the same sample. (**ii**) shows a closed ruffle containing PFF-gold. (iii) shows internalized PFF in a macropinosome. (**iv**) shows PFF in large forming MVBs in proximity to the membrane, suggesting PFF and membrane interaction. Scale bar= 200 nm for low magnification images, and 50 nm for insets. (**B**) In order to visualize the contents of lysosomes, which appear as highly electron dense compartments using EM, astrocytes exposed to PFF for 3-5 min were fixed and processed for EM; however, only *en bloc* uranyl acetate staining was conducted. Without staining of grids using uranyl acetate, PFF were easily visualized in multiple electron dense, lysosomal structures. Scale bar= 100 nm. (**C**) Uranyl acetate stained grids show highly electron dense lysosomes with PFF. Scale bar = 100 nm and 50 nm for insets.

Remarkably, gold-labelled PFFs are often found in close association with regions of membrane curvature as they begin to bud inward into electron-lucent organelles approximately 300-400 nm in diameter (**Fig. 6A**). Thus, PFF may contribute to the formation of MVBs. In fact, PFF are readily detected on the surface of vesicles in MVBs (**Fig. 6B**). MVBs, positive for LAMP1, containing PFF are also detected at the light level using STED microscopy and fluorescently-labeled PFF (**Fig. 6C**). Live-cell studies were performed to examine the dynamics of these structures, which appear to undergo multiple rounds of fusion and membrane budding (**Fig. 6D**/**E**). Thus, PFF enter cells via macropinocytosis and appear rapidly in lysosomes and MVBs.

**Figure 6.**
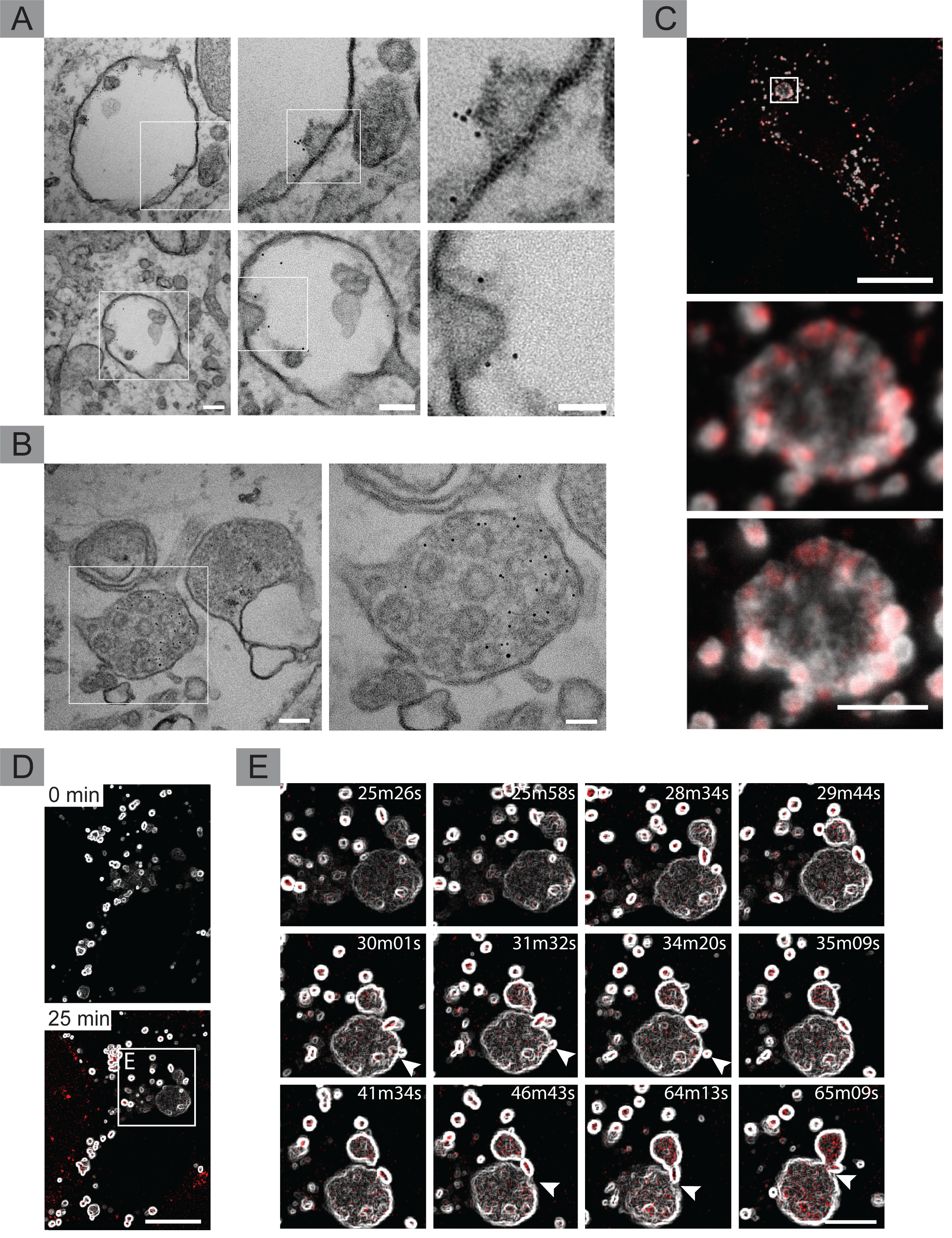
PFF trafficking to lysosomes and MVBs. (**A**) Astrocytes administered PFF for 10 min localize PFF in larger newly forming MVBs, always in proximity to the vesicular membrane, sometimes causing invaginations, signifying the early stages of exosome formation. Scale bar = 100 nm for low magnification, 80 nm for moderate magnification, 50 nm for highest magnification inset. (**B**) Localization of PFF outside of vesicles in electron dense MVBs signifies the progression of MVB maturation. Scale bar= 100 nm and 50 nm for inset. (**C**) Similar findings regarding the formation of MVBs (LAMP1 positive; white) containing PFF (red) was confirmed using STED microscopy. Scale bar = 20 µm for low magnification, and 2.5 µm for high magnification images. (**D**) Large-field image showing colocalization of PFF (red) with lysosomes(white): enhancement of lysosomal structures and their contents was carried out using “Find Edges” function in ImageJ, showing hollow lysosomal structures. Scale bar = 10 μm. (**E**) Higher magnification of images in D showing multiple budding events (arrowheads). Scale bar = 5 μm.

### PFF are transferred to naïve cells using exosomes

The spread of PFF throughout the nervous system requires that the fibrils escape cells and be transferred to neighboring naïve cells. Exosomes provide a mechanism for cell-to-cell transfer of protein, lipids, and other cellular molecules, and the presence of PFF in MVBs suggests a potential mechanism for cellular release. To test this hypothesis, U2OS cells with stable expression of CD63-GFP were incubated with fluorescently-labelled PFF for 24 h and then washed with trypsin and buffer before re-plating with PFF-naïve cells with stable expression of LAMP1-RFP. From 12-24 h following the start of co-culture, PFF were observed to transfer from the CD63-positive cells to the LAMP1-positive cells (**Fig. 7A**/**C**). Moreover, we also observed transfer of CD63 (**Fig. 7A**/**B**), suggesting that transfer of PFF involves exosomes. Similar results are seen with transfer to naïve cells expressing CD9-RFP (**Fig. S7A**). Notably, the transfer of both CD63 and PFF is blocked by the addition of manumycin A, a compound that disrupts exosome release (**Fig. 7A-C**).

**Figure 7.**
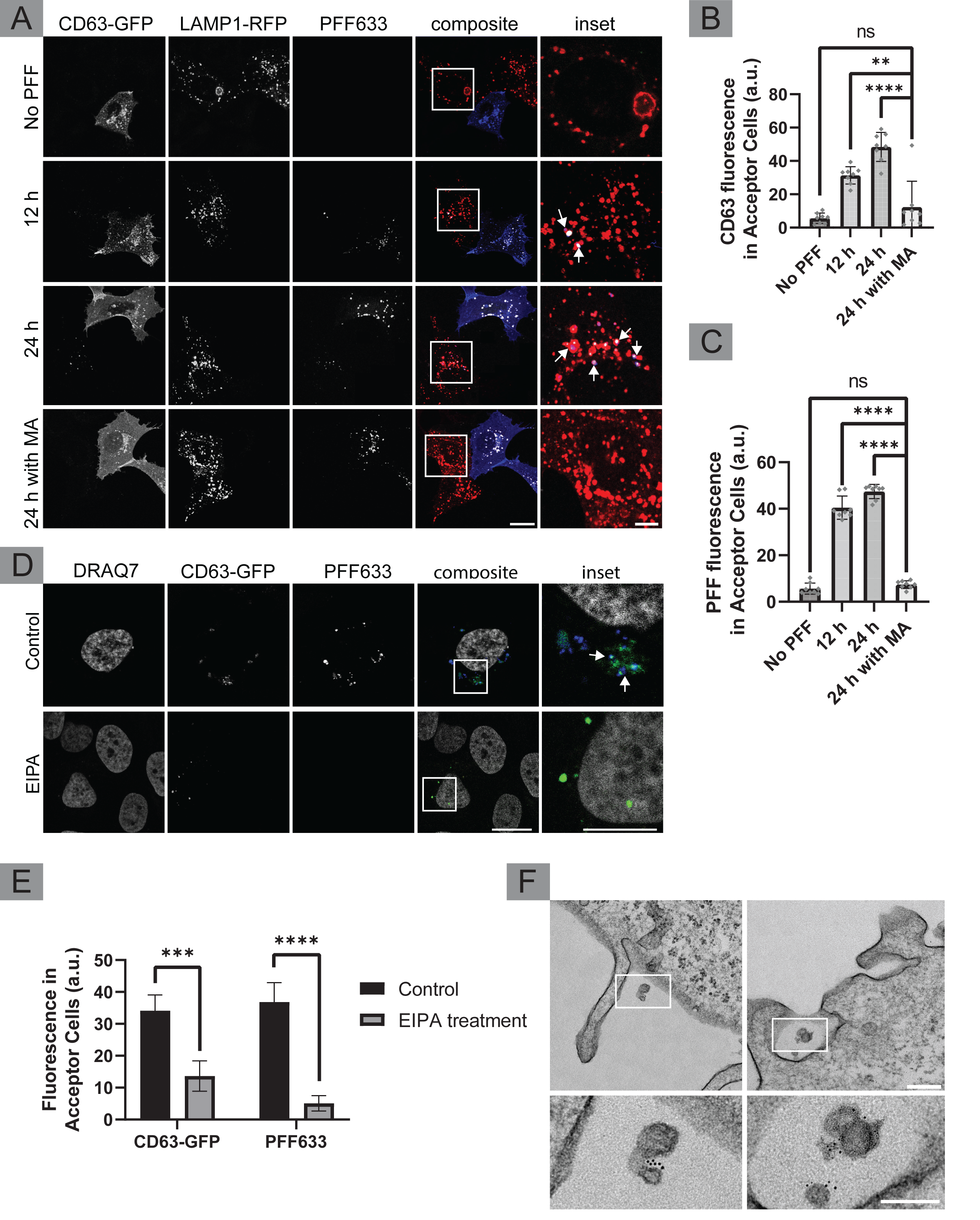
Exosomes containing PFF play a role in the transmission of PFF from donor to acceptor cells. (**A**) U2OS cells stably expressing CD63-GFP were exposed to PFF or PBS (vehicle control) for 24 h. CD63-GFP (donor cells; blue) were then trypsin washed three times, pelleted, trypsinized again, pelleted and PBS washed, prior to being co-plated with acceptor cells (PFF naïve cells stably expressing LAMP1-RFP; red). Donor and Acceptor cells were then incubated for 12 or 24 h. Half of the 24 h sample were given Manumycin A (MA) at 1.2 µM while the other half was given DMSO (vehicle control) at the time of plating. Both 12 h and 24 h samples show PFF (white) and CD63 (blue) fluorescence in acceptor cells, while the acceptor cells in the PBS and the MA condition show very little PFF and CD63 fluorescence. Arrows point to LAMP1-positive compartments in acceptor cells that contain both CD63 and PFF fluorescence. Scale bar = 10 µm for low magnification and 5 µm for insets. (**B**/**C**) Quantification of CD63 and PFF fluorescence in acceptor cells, respectively. n=8 for each condition (i.e., n = 32 total), from three independent experiments, mean ± SD. No PFF, 12 h and 24 h conditions were statistically compared (t-test) to the 24 h samples with MA; p < 0.0001 denoted as ****, p < 0.01 denoted as **. Individual data points are shown in grey. (**D**) At 24 h following the exposure of donor cells (CD63 positive) to PFF, cells were thoroughly trypsin washed and passaged onto plates and incubated with serum-free media for 36 h. The media was then collected, centrifuged for 5 min at 1000 RPM to pellet any floating cells, and given to acceptor cells (wildtype U2OS) some treated with DMSO (control) and others treated with EIPA. The cells were fixed following 6 h of incubation. Donor cells in control showed both PFF (blue) and CD63 (green) fluorescence, while EIPA treated cells did not. (**E**) Quantification of PFF and CD63 fluorescence in **D**. Multiple paired t-test was used to statistically compare CD63 and PFF fluorescence in acceptor cells (control vs EIPA); n = 6 per condition (n = 24 total) from three independent experiments; mean ± SD. p< 0.001 denoted as ***. (**F**) U2OS cells were given media (from donor cells treated with PFF-gold) isolated using the same protocol as **D** and incubated for 12 h. Cells were then processed for EM. Insets show extracellular vesicles with PFF-gold on their surface. Scale bar = 100 nm for low magnification images and 100 nm for inset.

We next took CD63 donor cells and incubated them with gold-labeled PFF for 24 h before extensive trypsin/buffer wash. The cells were then re-plated in serum-free media and incubated for 36 h before the media was collected, spun at low speeds to remove any cells or cell debris, and the supernatant transferred to naïve cells. We were able to capture images of extracellular vesicles in PFF naïve cells using confocal microscopy (**Fig. 7D**). We also found that the internalization of these exosomes was blocked by using EIPA (**Fig. 7E**). Following the same protocol to prepare EM samples, with the exception of a 48 h incubation time in fresh media supplemented with serum following re-plating, we found exosomes decorated with PFF in contact with naïve cells, sometimes seemingly near sites of membrane ruffles and forming macropinosomes (**Fig. 7F**). In investigating the response of cells to PFF, CD63-GFP expressing cells were given PFF for 24 h and were then washed with trypsin and re-plated. These cells often revealed CD63-positive ruffling and at the tip of ruffles, we could often spot PFF puncta colocalizing with CD63 (**Fig S7B**). We then isolated exosomes from WT U2OS cells that were exposed to PFF for 24 h. Exosomal isolation was performed using a series of centrifugation steps (Chhoy et al., 2021). Exosomes isolated from PFF treated and control cells showed fibril-like structures on their surface while the control exosomes did not (**Fig S7C**). Additionally, a portion of the exosomes isolated from PFF treated cells were trypsinized to determine whether PFF remains exclusively on the exosomal surface. Immunoblotting of the exosomal samples not only proved the identity of the exosomes (CD63-positive, GM130-negative) but additionally showed that the trypsinized exosomes did not contain any alpha-synuclein, hence PFF only reside on their surface (**Fig S7D**). Lastly, exosomes were isolated from U2OS cells that were given gold PFF for 24 h, trypsin washed, and then incubated for 48 h in fresh media (**Fig S7E**). Low centrifugation speeds allowed us to isolate exosomes, thanks to the nano-gold PFF, and allowed us to visualize them using EM. Gold covered exosomes were isolated in large clusters; however, we were able to show that PFF remains exclusively on the surface of exosomes. In summary, PFF that have entered cells by macropinocytosis, are trafficked to MVBs and appear to be delivered on exosomes to neighbouring cells.

## DISCUSSION

The mechanisms regarding the uptake of PFF remain incomplete (Bieri *et al*., 2018; Grozdanov and Danzer, 2018). Most studies investigating internalization of α-syn or PFF evaluate uptake hours following addition to cells (Desplats et al., 2009a; Hansen et al., 2011; Konno *et al*., 2012; Lee et al., 2008; Liu *et al*., 2007; Luk et al., 2016; Luk *et al*., 2009; Madeira *et al*., 2011; Rodriguez *et al*., 2018; Sung et al., 2001; Volpicelli-Daley *et al*., 2014; Volpicelli-Daley *et al*., 2011). While such uptake assays are valuable for genetic screening, they lack the temporal resolution to understand the pathways involved in endocytosis, an early event. We performed a detailed analysis of early events in endocytosis of PFF and were surprised to discover a unique uptake mechanism that allows the protein to reach lysosomes within 2 minutes, bypassing the early endosomal system.

Endocytosis of PFF has been thought to follow canonical CME pathways, entering cells via clathrin-coated pits and vesicles followed by trafficking through endosomes to lysosomes (Konno *et al*., 2012; Lee *et al*., 2008). However, we found no evidence for trafficking of PFF through early or recycling endosomes, inconsistent with a CME pathway. Moreover, using a previously established pool of CHC siRNA (Bayati et al., 2021; Galvez *et al*., 2007; Kim *et al*., 2011), we attained ∼95% knockdown efficiency, with no influence on PFF endocytosis. Thus, our data does not support a role for CME in uptake of PFF.

PFF induce the formation of actin- and Rac1-rich membrane ruffles, precursors of macropinocytic vesicles (Cox et al., 1997; Grimmer *et al*., 2002). Further, inhibitors of macropinocytosis (Commisso *et al*., 2014; Koivusalo *et al*., 2010; Zwartkruis and Burgering, 2013) including EIPA, LatA and LatB (Erami et al., 2017; Furstner et al., 2007; Morton et al., 2000; Wakatsuki et al., 2001) significantly inhibited uptake of PFF while not affecting EGF uptake. Our findings point to endocytosis of PFF through a form of macropinocytosis, where macropinosomes either mature into MVBs and lysosomes or fuse with pre-existing LAMP1-positive compartments. Recent papers show promising results regarding the role of actin in PFF internalization (Underwood et al., 2020; Zhang et al., 2020).

Aspects of our findings corroborate previous studies. First, although slower, previous research demonstrates the direct fusion of macropinosomes with lysosomes (Yoshida et al., 2018). Second, the findings of previous studies on dynamin inhibition and reduction in uptake of PFF can be due to dynamin’s role in some forms of macropinocytosis and its involvement in actin remodelling (Gu *et al*., 2010; Krueger et al., 2003; Mulherkar et al., 2011). In fact, at high concentrations, we observed that dynasore blocks internalization of PFF (data not shown).

Finally, amilorides, such as EIPA, which block uptake of PFF have been shown to have neuroprotective effects in PD (Arias et al., 2008).

Although previous studies attempted to examine the localization of exogenous α-syn using immuno-EM (Volpicelli-Daley *et al*., 2011), we conjugated PFF directly with gold. Many antibodies to α-syn cannot distinguish between different conformations of the protein (Kumar et al., 2020), making the direct conjugation of gold to PFF a more specific method. Consistent with previous literature (Vargas *et al*., 2017), we found that PFF were almost always in close association with membranes, whether at the cell surface, during internalization, or while in MVBs and lysosomes.

At early time points after addition to cells, we observed gold-PFF on inward invaginating vesicles in MVBs. At longer time points, gold-PFF were seen to be transported on exosomes, which confocal microscopy, demonstrated were CD63 positive. Two important discoveries were made as a result of this: first, PFF transmission is at least partly due to the exosomal transport of PFF; secondly, PFF reside on the surface of exosomes, rather than being contained in the lumen of exosomal vesicles. Further, exosomes isolated from cells exposed to PFF contained PFF on their outer surface. Biochemically, we were able to show that PFF can be removed from exosomes through trypsinization; this would only be possible if PFF resided on the surface of exosomes. We then used Manumycin A, a drug that blocks the release of CD63-positive exosomes (Datta et al., 2017), and found that Manumycin A disrupts PFF transmission from cell-to-cell. We suspect that since PFF resides on the outside of exosomes, that drugs like EIPA, LatA, and LatB that blocked its initial uptake, can be used to block PFF spread from cell-to-cell. Our hypothesis was confirmed with our experiment using EIPA, showing the inhibition of exosomal uptake by PFF-naïve cells.

Although all evidence presented indicates that PFF use macropinocytosis to gain access into cells, we are also cognizant that the properties of α-syn itself could drive the membrane curvature and protrusions. As previously observed, PFF associate and may even drive membrane curvature (Vargas *et al*., 2017; Westphal and Chandra, 2013). This could mean that PFF drives its own internalization into the cell by causing membrane curvature, and then, once trafficked to MVBs/lysosomes, it drives membrane invaginations, resulting in exosomal formation and eventually its release. Once released, PFF then goes on to disrupt other cells, all the while remaining on the outside of exosomes, allowing it to drive membrane curvature that would drive its internalization in neighboring cells. Although PFF is only an oligomer, it certainly has many prion-like characteristics. Above all else, PFF may be taking advantage of the α-syn’s interaction with membranes to drive its own endocytosis, exocytosis, and drive the production of more aggregated α-syn. It is this evolutionary drive for PFF to create more fibrillated forms of α-syn, that truly makes it more than just an oligomer, and more like a prion.

In conclusion, our data demonstrate that PFF enter cells via macropinocytosis, with seemingly direct transfer to MVB and lysosomes, bypassing early endosomal pathways. It remains unclear if this represents fusion of macropinosomes with MVB and lysosomes or maturation of macropinosomes into these structures. A portion of the PFF that enter the cells are subsequently secreted on exosomes, providing a mechanism for cell-to-cell transport, and yet our data does not explain how fibrillar αSyn interacts with αSyn in the cytosol to allow prion-like propagation. Regardless, inhibition of macropinocytic pathways may prove useful in limiting the progression of PD and other synucleinopathies.

## ACKNOWLEDGEMENTS

We acknowledge the Neuro Microscopy Imaging Centre and Advanced BioImaging Facility and the Facility for Electron Microscopy Research at McGill University. We thank Dr. Michael Davidson and Dr. Paul Luzio for the LAMP1 and CD63 plasmids, respectively. We also thank Drs. Sabatini and Zoncu for the HA-TMEM192 plasmids. AB is supported by Fonds de recherche du Québec doctoral award and a studentship from the Parkinson Society of Canada. This work was supported by a grant from the Canada First Research Excellence Fund, Healthy Brain, Healthy Lives, McGill University, awarded to PSM. PSM is a Distinguished James McGill Professor and Fellow of the Royal Society of Canada.

## AUTHOR CONTRIBUTIONS

A.B. planned and conducted the experiments and wrote the manuscript with P.S.M. E.B. performed lysosomal immunoprecipitation experiment along with analyzing protein expression through western blotting. C.H. provided NPCs and differentiated iPSCs into cortical and dopaminergic neurons. W.L. produced and characterized PFFs. W.E.R. conducted 24 h PFF uptake experiment with dopaminergic NPCs using LatA. C.Z. provided us with additional NPCs and differentiated iPSCs. R.S. measured and analyzed PFF length from multiple batches of PFF using dynamic light scattering. E.D.C.P. characterized and measured PFF size via EM. B.V. suggested experiments investigating macropinocytosis. H.M.M., E.A.F. and T.M.D. edited the manuscript and helped provide overall direction of the project. P.S.M. funded and supervised the project, aided A.B. in planning the experiments, and wrote the manuscript with A.B.

## STAR METHODS

### KEY RESOURCE TABLE

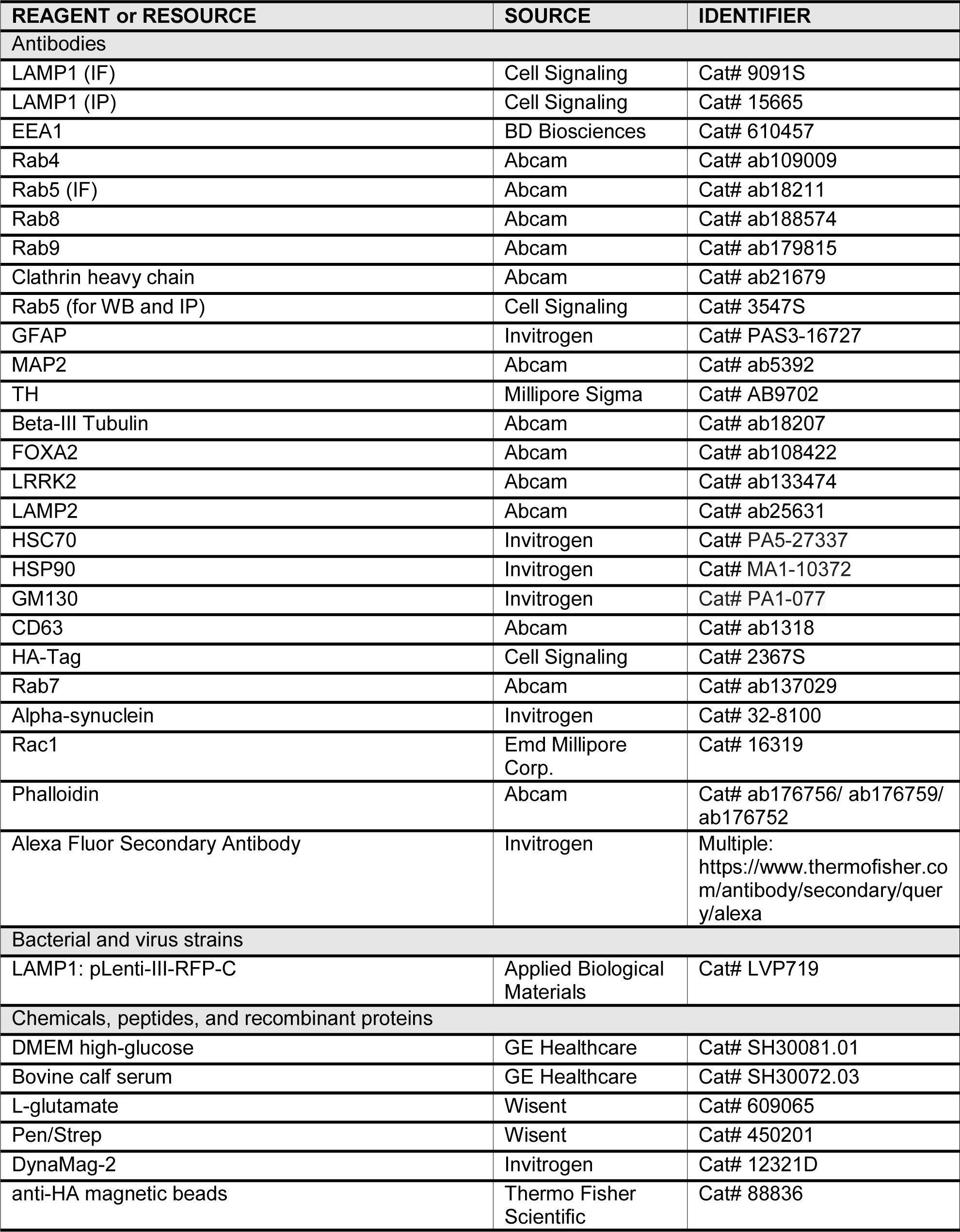
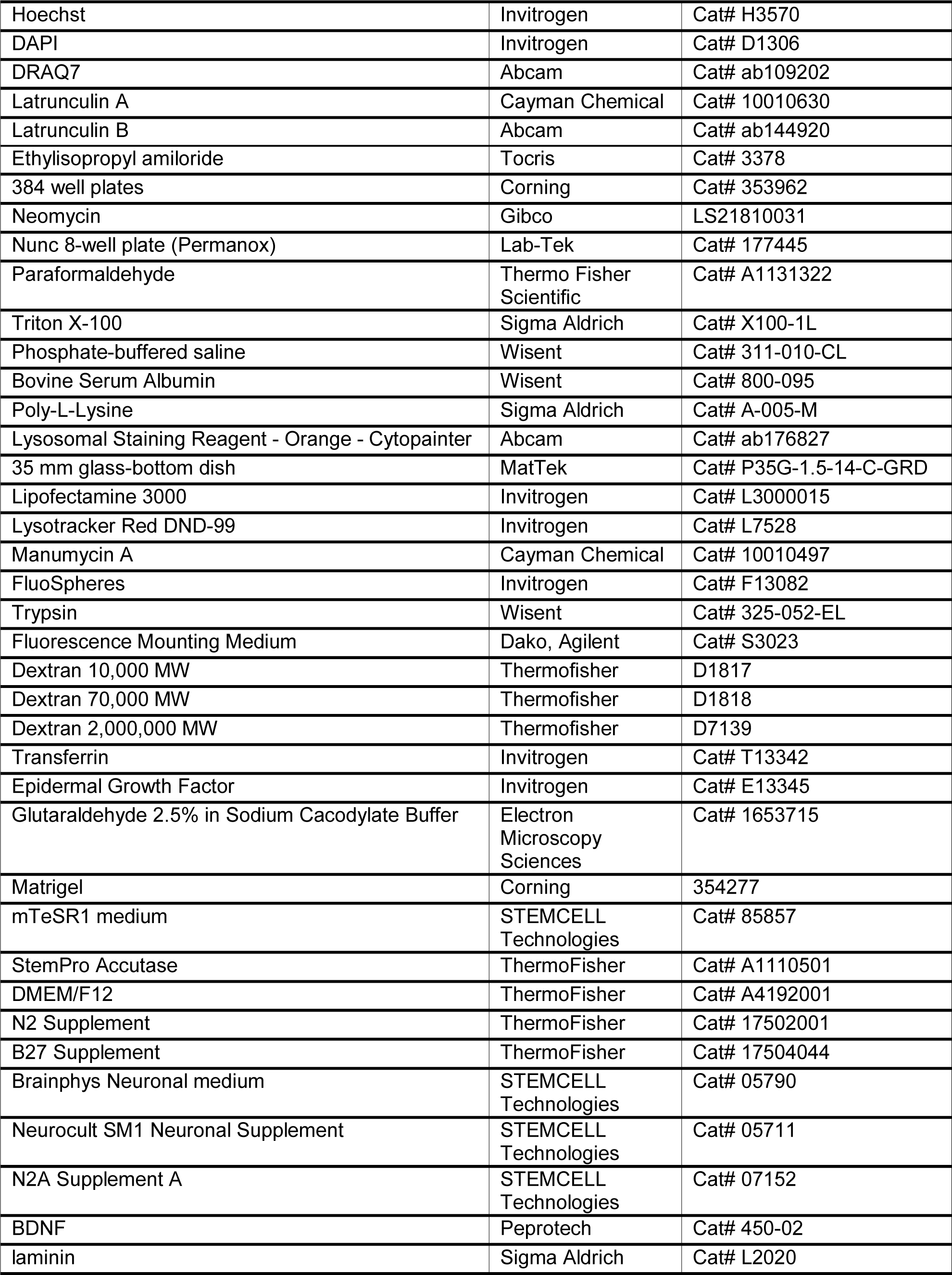
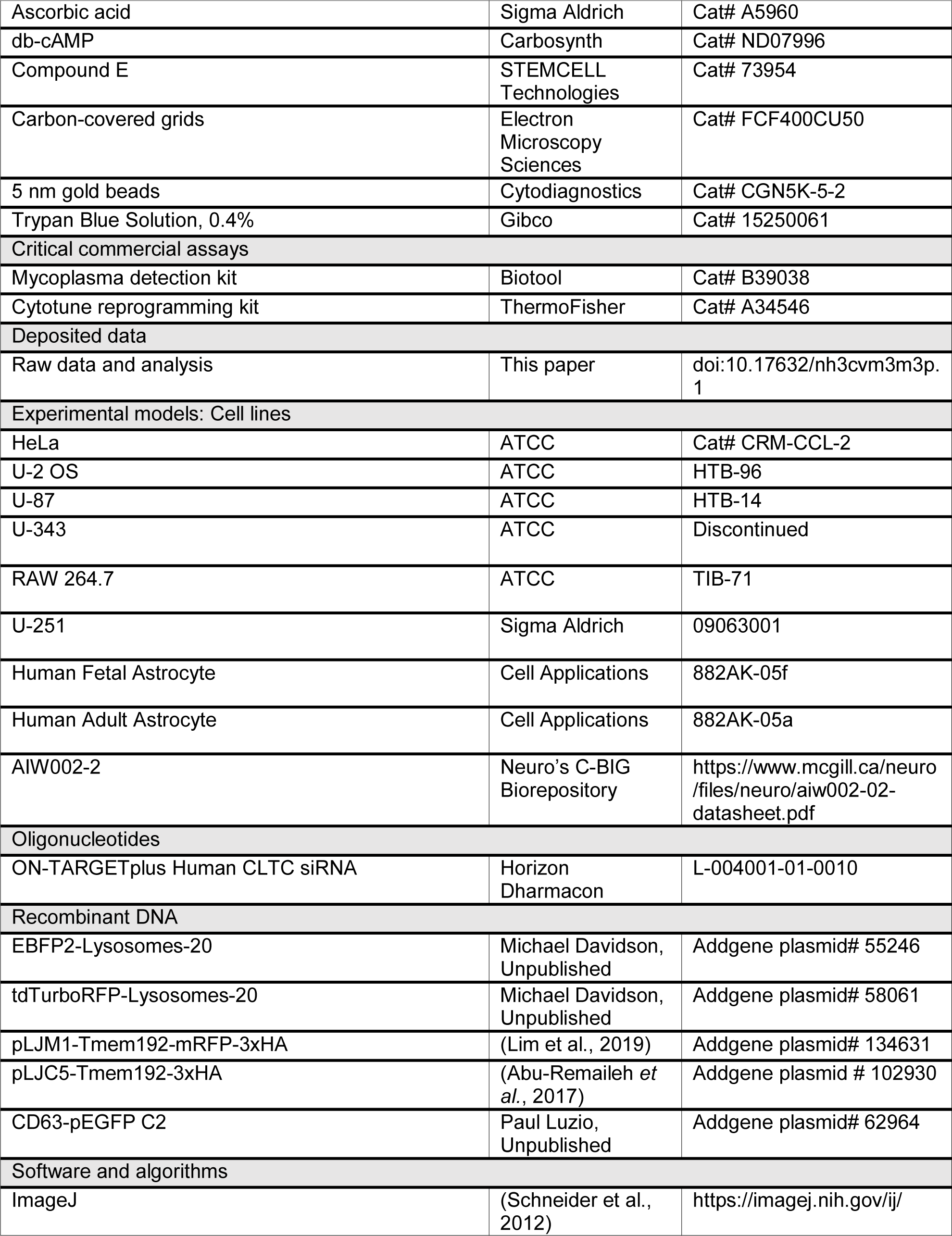
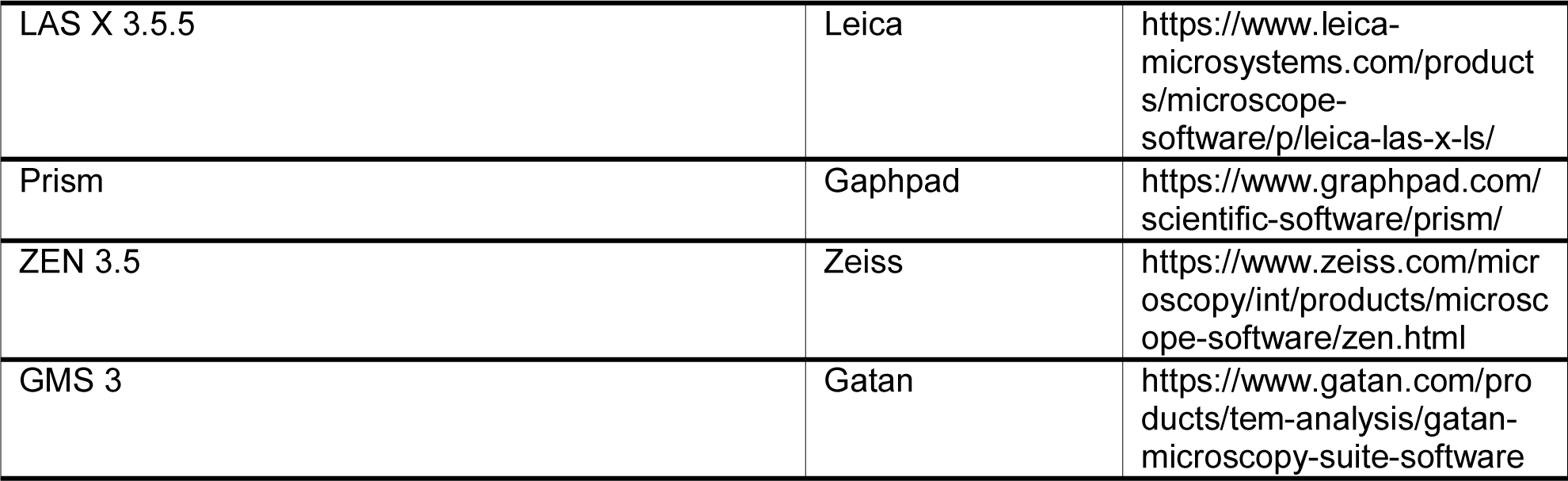

### RESOURCE AVAILABILITY

#### Lead Contact

Further information and requests for resources and reagents should be directed to the Lead Contact, Dr. Peter Scott McPherson and will be fulfilled.

#### Materials Availability

This study did not generate new unique reagents.

#### Data and Code Availability

- All the raw data, along with statistical calculations used in this paper have been submitted in a spreadsheet file along with the manuscript to Cell Press. All data reported in this paper will be shared by the lead contact upon request.
- This paper does not report original code.
- Any additional information required to reanalyze the data reported in this paper is available from the lead contact upon request.

### EXPERIMENTAL MODELS AND SUBJECT DETAILS

#### Cell lines

HeLa, U2OS, U87, U-343, RAW 264.7 were obtained from American Type Culture Collection (Cat# CRM-CCL-2, HTB-96, HTB-14, discontinued, TIB-71, respectively). U-251 cells were obtained from Sigma (Cat# 09063001). Human fetal and adult astrocytes were obtained from Cell Applications (Cat# 882AK-05f and 882AK-05a, respectively). For studies with iPSCs, we used the line AIW002-2 obtained from the Neuro’s C-BIG Biorepository. This line was reprogrammed from peripheral blood mononuclear cells of a healthy donor with the Cytotune reprogramming kit (ThermoFisher, Cat# A34546). The process of reprogramming and quality control profiling for this iPSC was outlined in a previous study (Chen et al., 2021). The use of iPSCs in this project is approved by the McGill University Health Centre Research Ethics Board (DURCAN_IPSC / 2019-5374).

All cells were cultured in DMEM high-glucose (GE Healthcare, Cat# SH30081.01) containing 10% bovine calf serum (GE Healthcare, Cat# SH30072.03), 2 mM L-glutamate (Wisent, Cat# 609065, 100 IU penicillin and 100 μg/ml streptomycin (Wisent cat# 450201). Cell lines were routinely checked for mycoplasma contamination using the mycoplasma detection kit (Biotool cat# B39038).

#### Production, Characterization, and Nano-Gold labelling of PFF

Production and characterization of recombinant α-syn monomers and PFF have been described previously (Del Cid Pellitero *et al*., 2019; Maneca *et al*., 2019). Both electron microscopy and dynamic light scattering were used for the characterization of α-syn monomers and PFF (**Fig. S1A-B**). Previously characterized PFF, was then conjugated with 5 nm gold beads (Cytodiagnostics, Cat# CGN5K-5-2), immediately before experimental use. Cytodianostic’s conjugation protocol was used to conjugate gold onto PFF. Following conjugation, some PFF was collected for characterization on carbon-covered grids (Electron Microscopy Sciences, Cat# FCF400CU50) (**Fig. S1C**).

#### IPSC Culturing

AIW002-2 hiPSC cultures were maintained as feeder-free cultures following the protocol described previously (Chen *et al*., 2021). AIW002-2 hiPSCs were plated onto Matrigel (Corning, Cat# 354277)-coated plates containing mTeSR1 medium (Stemcell Technologies, Cat# 85857). The culture medium was changed daily until cells reached ∼80% or required confluency (usually 5-7 days after plating). The cells were then passaged, frozen, or differentiated. A previously described protocol was used to generate ventral midbrain dopaminergic neural progenitor cells (Jefri et al., 2020). Dopaminergic neural progenitor cells were dissociated with StemPro Accutase Cell Dissociation Reagent (ThermoFisher, Cat# A1110501) into single-cell suspensions. 50,000 cells were plated onto coated coverslips in 24-well plates with neural progenitor plating medium (DMEM/F12 supplemented with N2, B27 supplement; ThermoFisher, Cat# A4192001, 17502001, 17504044). To further differentiate into dopaminergic neurons, neural progenitor medium was switched to dopaminergic neural differentiation medium (Brainphys Neuronal medium, STEMCELL Technologies, Cat# 05790) supplemented with N2A Supplement A (STEMCELL Technologies; Cat# 07152), Neurocult SM1 Neuronal Supplement (STEMCELL Technologies; Cat# 05711), BDNF (20 ng/mL; Peprotech, Cat# 450-02), GDNF (20 ng/mL; Peprotech, Cat# 450-10), Compound E (0.1 μM; STEMCELL Technologies, Cat# 73954), db-cAMP (0.5 mM; Carbosynth, Cat# ND07996), Ascorbic acid (200 μM; Sigma Aldrich, Cat# A5960) and laminin (1 μg/mL, Sigma Aldrich, Cat# L2020).

#### Plasmids and lentivirus

EBFP2-Lysosomes-20, tdTurboRFP-Lysosomes-20 were gifts from Michael Davidson (Addgene plasmid# 55246 and 58061). pLJM1-Tmem192-mRFP-3xHA was a gift from Roberto Zoncu (Addgene, plasmid# 134631). pLJC5-Tmem192-3xHA was a gift from David Sabatini (Addgene plasmid # 102930). CD63-pEGFP C2 was a gift from Paul Luzio (Addgene, Cat# 62964). LAMP1-RFP lentivirus was obtained from Applied Biological Materials Inc. (Cat# LVP719).

### METHOD DETAILS

#### Trypan Blue Exclusion Assay

HeLa (ATCC, Cat# CRM-CCL-2) cells were grown on poly-L-lysine coated 35 mm glass-bottom dishes (MatTek, Cat# P35G-1.5-14-C-GRD) for 48 h to 50% confluency. When ready for imaging, cells were stained with Hoescht (Invitrogen, Cat# H3570) for 30 min and dishes were placed in the pre-heated live imaging chamber of the Zeiss LSM-880 confocal microscope.

Imaging was commenced before the addition of PFF tagged with Alexa Fluor 488 (PFF488) at a speed of 1 frame/sec. PFF was added during live imaging.

#### PFF Live Internalization Assay

U2OS (ATCC, HTB-96) cells were grown on Poly-L-Lysine (Sigma Aldrich, Cat# A-005-M) coated 35 mm glass-bottom dish (MatTek, Cat# P35G-1.5-14-C-GRD) for 72 h to 70% confluency. Before imaging, cells were stained with Lysosomal Cytopainter (Abcam, Cat# ab176827) for 30min. Dishes were placed in the pre-heated live imaging chamber of the Zeiss LSM-880 confocal microscope. Imaging was commenced before the addition of PFF at a speed of 1 frame/sec using the Airy imaging mode for higher resolution (∼1.3x resolution of conventional confocal microscopy). PFF488 was added during imaging.

#### Live MVB Assay

These live experiments were carried out the same way as above. For the hollow lysosome morphology, the “Find Edges” processing was used on ImageJ (NIH, https://imagej.net/software/fiji/).

#### Lysosomal Staining in Fixed Samples

Lysosomal staining was achieved in one of the following ways: transfection with plasmids stated above with the aid of Lipofectamine 3000 (ThermoFisher, Invitrogen, Cat# L3000015), fixable Lysotracker (ThermoFisher, Invitrogen, Cat# L7528) for staining of lysosomes in neurons and neural progenitor cells, or with the use of LAMP1 antibody (Cell Signaling, Cat# 9091S. Transfection was done 24 h prior to experimentation. Staining with Lysotracker was done 30 min before experimentation. LAMP1 antibody staining was done following fixation and permeabilization.

#### PFF Endocytosis Assays

Cells mounted on coverslips were plated in 24 well plates at 37°C with 200 µl of media in each well. Immediately prior to experimentation, cells were taken out of incubators and placed in the cell culture hood. PFF aliquots were then removed from dry ice, diluted with serum-free media, and added to each well. Cells were then incubated at 37°C for 0, 2, 10, 30, or longer. Cells were then placed on ice and washed with trypsin (Wisent, Cat# 325-052-EL) for 90 sec (to remove extracellular PFF). Following 2 washes with PBS (Wisent, Cat# 311-010-CL), cells were fixed. In experiments where trypsin was not used, cells were washed three times with PBS and fixed. In experiments using iPSC neurons, trypsinization was avoided and only PBS was used to wash cells. Same protocol was followed for samples prepared for EM, except for plating cells on Nunc 8-well dishes (Lab-Tek, Nunc, Thermo Scientific, Cat# 177445) and fixation with Glutaraldehyde and not PFA.

#### PFF Long-term Endocytosis Assays

Cell plating and culturing was done as described above. Cells were then incubated at 37°C for 24 h. Following 24 h, cells were trypsin washed three times, pelleted, resuspended and re-plated onto fresh coverslips. For iPSC neurons, trypsinization (Wisent, Cat# 325-052-EL) and re-plating was avoided and only the media was changed at 24 h. Cells were then incubated for varying amount time afterwards. Prior to imaging, sample fixation and mounting were conducted as above.

For quantifying the PFF fluorescence over 14 d, the same protocol as above was employed; however, in order to control for cell growth, human adult astrocytes (Cell Applications, Cat# 882AK-05a) at passage 7 were used as these cells have a very slow doubling time. This allowed us to control for growth without needing to inhibit cell replication through treatment with chemicals that could have a negative affect on PFF uptake, fluorescence, or retention.

#### Macropinocytic Inhibitors

Ethylisopropyl amiloride (EIPA; R&D Systems, Tocris, Cat# 3378), Latrunculin A (LatA; Cayman Chemical, Cat# 10010630), and Latrunculin B (LatB, respectively; abcam, Cat# ab144920, ab144291) were the macropinocytic inhibitors used in this study. Cells normally incubated in media with serum were washed 3 times with serum-free media to remove any remaining serum. Serum-free media was added with a final concentration of either 20 µM of EIPA, 0-2 µM of LatA, 5 µM of LatB. Cells were then incubated at 37^°^C for 30 min before experimentation for EIPA and LatB, and 1 h for LatA. Due to the autofluorescence of EIPA, DRAQ7 (abcam, Cat# ab109202) was used to stain cell nuclei.

#### LatA Inhibition in Dopaminergic NPCs

Dopaminergic NPCs were seeded in polyornithin/ laminin coated 384 well plates (Corning, Cat# 353962) at 4000cells/ well. After 24 h Latrunculin A (LatA; Cayman Chemical, Cat# 10010630) was added. After 1 h, Alexa488-PFF was added. After another 24 h incubation period, the wells were washed twice with PBS (Wisent, Cat# 311-010-CL) and the cells fixed with 2% FA/PBS.

Cells were counterstained with Hoechst 33342 (ThermoFisher, Cat# H3570) and imaged on a high content imager (CellInsight CX7, ThermoFisher Scientific). The amount of intracellular Alexa488-PFF was measured as the total intensity of Alexa488 fluorescent stain in the perinuclear area. Nuclei count and nuclear area (px^2^) were also obtained as indicators for cell toxicity. All data was normalized against DMSO (vehicle) only controls. Data represents the mean and standard deviation of 3 independent experiments.

#### Lysosomal Immunoprecipitation

U2OS cells stably expressing 3xHA-TMEM192-RFP were incubated with PFF for varying lengths of time, washed with trypsin (Wisent, Cat# 325-052-EL) three times to remove extracellular PFF, and then trypsinized and pelleted. Lysosomes were immunoprecipitated based on the protocol described in Abu-Remaileh et al. (2018). Briefly, cells were washed twice with KPBS and centrifuged at 4°C for 2 minutes at 1000 x g. Pelleted cells were resuspended in lysis buffer (20 mM HEPES pH 7.4, 100 mM NaCl, and protease and phosphatase inhibitor cocktail) and manually homogenized with 25 up and down strokes. Homogenates were centrifuged for 2 min at 4°C at 1000 x g and 10% of the total volume for each IP was reserved for starting material (SM) fractions. Homogenates were incubated with gentle rocking at 4°C for 3 minutes with 100 μL anti-HA magnetic beads (ThermoFisher, Thermo Scientific, Cat# 88836) pre-washed with KPBS. After collecting the total IP volume for the unbound materials (UM), immunoprecipitates were washed three times with KPBS using a DynaMag-2 Magnet (ThermoFisher, Invitrogen, Cat# 12321D) before resuspension in 1X SDS-PAGE sample buffer. Protein aliquots were then analyzed by SDS-PAGE and Immunoblot.

For samples to be processed via confocal microscopy, 20 μL pre-washed anti-HA beads were used in the IP and were resuspended in using PFA to resuspend and fix them. The sample was then pelleted and resuspended with PBS (Wisent, Cat# 311-010-CL) and mounted onto slides.

#### PFF Intercellular (with Contact) Transfer Assay

U2OS cells were transfected with CD63-EGFP or LAMP1-tdTurboRFP using Lipofectamine 3000 transfection reagent and incubated for 24 h. Cell media was then changed, and cell selection process was initiated through the addition of Neomycin (Gibco cat# LS21810031). U2OS cells stably expressing CD63-GFP were exposed to PFF or PBS (vehicle control) for 24 h. CD63-GFP (donor cells) were then trypsin washed three times, pelleted, trypsinized again, pelleted and PBS washed, prior to being co-plated with acceptor cells (PFF naïve cells stably expressing LAMP1-tdTurboRFP). Donor and Acceptor cells were then incubated for 12 or 24 h. Half of the 24 h sample were given Manumycin A (MA; Cayman Chemical, Cat# 10010497) at 1.2 µM while the other half was given DMSO (vehicle control) at the time of plating. Cells were then fixed, mounted and imaged.

#### PFF Intercellular (without Contact) Transfer Assay

U2OS cells were stably expressing with CD63-EGFP, the cells were exposed to PFF, for 24 h then trypsin washed three times and re-plated prior to the addition of serum-free media. The media was collected following 36 h of incubation at 37°C and given to naïve U2OS cells. Some of the cells given this media were exposed to EIPA while others were given DMSO as a vehicle control. For fluorescent microscopy samples, the naïve cells were plated on coverslips, while for samples to be examined with EM, naïve cells were plated on Nunc 8-well plates (Lab-Tek, Nunc, Thermo Scientific, Cat# 177445).

#### Fixation and Antibody Staining Following PFF Uptake

Fixation was done with 2% freshly made paraformaldehyde (PFA; Thermo Fisher Scientific, A1131322) for 10-15 min on ice. In experiments where antibody staining was done, cells were then blocked and permeabilized for 30 min using 0.05% Triton X-100 (Sigma, Cat# X100-1L) in Phosphate-buffered saline (PBS; Wisent, Cat# 311-010-CL) along with 5% BSA (Wisent, Cat# 800-095). Coverslips were then transferred into a wet chamber and incubated with 1:500 dilution of the antibody in 0.01% Triton X-100 and 5% BSA. Cells were incubated with the diluted antibody for 2 h at room temperature. Coverslips were then gently washed 3 times with PBS, and 1:500 dilution of secondary antibody was added in 0.01% Triton X-100 and 5% BSA. Cells were then washed 2 more times with PBS and stained with DAPI (ThermoFisher, Invitrogen, Cat# D1306) for 10 min at 1 µg/ml concentration. Coverslips were then mounted on a glass slide using Fluorescence Mounting Medium (Dako, Agilent, Cat#S3023). All fixed samples were then imaged using a Leica TCS SP8 confocal microscope. STED samples were imaged using Abberior STED super-resolution nanoscope.

We initially found that PFF staining was severely affected following permeabilization limiting our ability to use antibodies in our experiments. We found that in cells with higher uptake of PFF (e.g., astrocytes), we could permeabilize samples and still retain some PFF fluorescence. We also found that instead of fixation and permeabilization with methanol or PFA and high concentrations of Triton X-100 (i.e., 0.1-0.5% Triton volume/volume), that a softer fixation with 2% PFA and low concentration of Triton X-100 for permeabilization (i.e., 0.05% Triton) allowed for the preservation of PFF fluorescence. Following our understanding of how to mediate the deleterious effects of permeabilization on PFF fluorescence in fixed samples, we could then employ antibodies in our experiments to stain for endosomal markers.

#### CHC KD

U2OS cells at 60% confluency were transfected with siRNA retrieved from Dharmacon (SMARTpool, ONTARGETplus, see KRT) or control siRNA (Dharmacon; ON-TARGETplus CONTROL) using Lipofectamine 3000 (ThermoFisher, Invitrogen, Cat# L3000015). At 24 h following transfection, cells were passaged, with some cells retrieved for imaging experiments and mounted onto coverslips. At 48 h, cells were collected in HEPES lysis buffer (20 mM HEPES, 150 mM sodium chloride, 1% Triton X-100, pH 7.4) accompanied with protease inhibitors. Cells in lysis buffer were then placed at 4°C and gently rocked for 30 min. Lysates were then spun at 238,700 x g for 15 min at 4°C. Lysates were run on 5-16% gradient polyacrylamide gel and transferred onto nitrocellulose membranes. Subsequently, proteins were visualized using Ponceau staining. Blots were then blocked with 5% milk for 1 h. Antibodies were then incubated overnight at 4% in bovine serum albumin/TBS/0.1 Tween 20. The secondary antibody was incubated at 1:2500 dilution in 5% milk/TBS/0.1 Tween 20 for 1 h at room temperature. Concurrently, cells mounted onto coverslips underwent the endocytosis assay with both knockdown and control cells mounted onto the same coverslips.

#### Fluorosphere and Dextran Uptake Assays

RAW 264.7 cells (ATCC, Cat# TIB-71) were plated onto coverslips treated with Poly-L-Lysine (Sigma, Cat# A-005-M). FluoSpheres (ThermoFisher, Invitrogen, Cat# F13082), latex beads of 1 µm diameter, and orange fluorescence were incubated with and without PFF (before and simultaneously). Cells were then washed with trypsin (Wisent, Cat# 325-052-EL), washed twice with PFF, and fixed. Cells were then counterstained with DAPI prior to mounting using Fluorescence Mounting Medium (Dako, Agilent, Cat#S3023).

U2OS cells were used in the dextran and PFF co-uptake experiment. U2OS cells were plated and prepared as described above. Three different dextran (10,000 MW, 70,000 MW, 2,000,000 MW; ThermoFisher, Cat# D1817, D1818, D7139) were separately added to cells prior or with PFF addition.

#### Membrane Ruffling Assay

Cellular response to PFF addition was done using human fetal astrocytes (Cell Applications, 882AK-05f). Cells were treated with PFF, and their recruitment of Rac1 (Emd Millipore Corporation, Cat# 16319) to the cell surface and colocalization with F-actin (phalloidin; Abcam, Cat# A22287) was analyzed. For negative and positive control, Transferrin (Tf; ThermoFisher, Invitrogen, Cat# T13342) Epidermal growth factor (EGF; ThermoFisher, Invitrogen, Cat# E13345) were added to cells, respectively. A null condition was also in place, were cells were administered PBS (a vehicle control for PFF).

#### Exosomal Isolation

For exosomal isolation, a previously published protocol was followed (Chhoy *et al*., 2021). Following isolation, samples were prepared for immunoblotting and EM. For immunoblotting, a portion of exosomes retrieved from PFF treated cells were treated with trypsin for 5 min. The exosomes were then re-pelleted at 10,000 g’s, and the supernatant removed. These trypsinized exosomes were run alongside exosomes collected from PBS and PFF treated cells, to compare PFF content. For isolation of exosomes collected from cells exposed to PFF-gold, a single centrifugation step at 10,000 g’s was used to pellet exosomes.

#### TEM

Human astrocytes and U2OS cells were plated onto 8 well chamber slides (Lab-Tek, Nunc, Thermo Scientific, Cat# 177445) and were administered PFF conjugated with gold. Cells were then fixed with glutaraldehyde 2.5% in 0.1M sodium cacodylate buffer (Electron Microscopy, Cat# 1653715), post-fixed with 1% OsO_4_ and 1.5% potassium ferrocyanide in sodium cacodylate buffer. Cells were then *en bloc* stained with 4% uranyl acetate. Post-embedding, some grids were stained with uranyl acetate for enhanced membrane staining. Samples were viewed with a Tecnai Spirit 120 kV electron microscope and captured using a Gatan Ultrascan 4000 camera.

#### Graphical Abstract

Created with BioRender.com

### QUANTIFICATION AND STATISTICAL ANALYSIS

#### Colocalization Rate Calculation

Colocalization was quantified using the Leica LAS X software. A 50% background threshold was set to eliminate all background fluorescence and fluorescent bleed-through across channels. Colocalization rate was used for graphs and statistical calculations. Colocalization was calculated by measuring the colocalization area between two channels (in µm^2^) and dividing it by the foreground area (foreground area = image area divided by total area). This allows us to look at what portion of the image is taken up by colocalization, a great indicator of the degree of colocalization between two channels, while considering the area that each channel occupies.

This is based on MOC, mander’s overlapp coefficient.

#### Quantification and Statistics

For all quantifications, including uptake and colocalization, the Leica LAS X and ImageJ (imagej.net) software were used. Readout from the LAS X software included Pearson Correlation Coefficient and colocalization rate expressed in percentage. In each experiment, images were selected from a large field imaged at low quality using the Leica SP8 spiral scan. From the large field, regions were selected that contained a minimum of 4 cells. Graphs were then prepared using GraphPad Prism 9 software. For statistical comparisons, Student’s t-test (unpaired) and one-way ANOVA were used. When significance was detected under ANOVA, multiple comparisons Tukey’s test was conducted to find means that are significantly different across conditions. All data is shown as mean ± SD when quantifying uptake and mean ± minimum and maximum values for colocalization. For statistical significance, p < 0.05 was used.

## REFERENCES

1. Abu-Remaileh, M., Wyant, G.A., Kim, C., Laqtom, N.N., Abbasi, M., Chan, S.H., Freinkman, E., and Sabatini, D.M. (2017). Lysosomal metabolomics reveals V-ATPase- and mTOR-dependent regulation of amino acid efflux from lysosomes. Science 358, 807–813. 10.1126/science.aan6298.

2. Arias, R.L., Sung, M.L., Vasylyev, D., Zhang, M.Y., Albinson, K., Kubek, K., Kagan, N., Beyer, C., Lin, Q., Dwyer, J.M., et al. (2008). Amiloride is neuroprotective in an MPTP model of Parkinson’s disease. Neurobiol Dis 31, 334–341. 10.1016/j.nbd.2008.05.008.

3. Bayati, A., Kumar, R., Francis, V., and McPherson, P.S. (2021). SARS-CoV-2 infects cells following viral entry via clathrin-mediated endocytosis. J Biol Chem, 100306. 10.1016/j.jbc.2021.100306.

4. Betemps, D., Verchere, J., Brot, S., Morignat, E., Bousset, L., Gaillard, D., Lakhdar, L., Melki, R., and Baron, T. (2014). Alpha-synuclein spreading in M83 mice brain revealed by detection of pathological alpha-synuclein by enhanced ELISA. Acta Neuropathol Commun 2, 29. 10.1186/2051-5960-2-29.

5. Bieri, G., Gitler, A.D., and Brahic, M. (2018). Internalization, axonal transport and release of fibrillar forms of alpha-synuclein. Neurobiol Dis 109, 219–225. 10.1016/j.nbd.2017.03.007.

6. Braak, H., Tredici, K.D., Rüb, U., de Vos, R.A.I., Jansen Steur, E.N.H., and Braak, E. (2003). Staging of brain pathology related to sporadic Parkinson’s disease. Neurobiology of Aging 24, 197–211. 10.1016/s0197-4580(02)00065-9.

7. Bryant, D.M., Kerr, M.C., Hammond, L.A., Joseph, S.R., Mostov, K.E., Teasdale, R.D., and Stow, J.L. (2007). EGF induces macropinocytosis and SNX1-modulated recycling of E-cadherin. J Cell Sci 120, 1818–1828. 10.1242/jcs.000653.

8. Chen, C.X.-Q., Abdian, N., Maussion, G., Thomas, R.A., Demirova, I., Cai, E., Tabatabaei, M., Beitel, L.K., Karamchandani, J., Fon, E.A., and Durcan, T.M. (2021). Standardized quality control workflow to evaluate the reproducibility and differentiation potential of human iPSCs into neurons. bioRxiv, 2021.2001.2013.426620. 10.1101/2021.01.13.426620.

9. Chhoy, P., Brown, C.W., Amante, J.J., and Mercurio, A.M. (2021). Protocol for the separation of extracellular vesicles by ultracentrifugation from in vitro cell culture models. STAR Protoc 2, 100303. 10.1016/j.xpro.2021.100303.

10. Commisso, C., Flinn, R.J., and Bar-Sagi, D. (2014). Determining the macropinocytic index of cells through a quantitative image-based assay. Nat Protoc 9, 182–192. 10.1038/nprot.2014.004.

11. Condon, N.D., Heddleston, J.M., Chew, T.L., Luo, L., McPherson, P.S., Ioannou, M.S., Hodgson, L., Stow, J.L., and Wall, A.A. (2018). Macropinosome formation by tent pole ruffling in macrophages. J Cell Biol 217, 3873–3885. 10.1083/jcb.201804137.

12. Conway, K.A., Harper, J.D., and Lansbury, P.T. (1998). Accelerated in vitro fibril formation by a mutant alpha-synuclein linked to early-onset Parkinson disease. Nat Med 4, 1318–1320. 10.1038/3311.

13. Conway, K.A., Lee, S.J., Rochet, J.C., Ding, T.T., Williamson, R.E., and Lansbury, P.T., Jr. (2000). Acceleration of oligomerization, not fibrillization, is a shared property of both alpha-synuclein mutations linked to early-onset Parkinson’s disease: implications for pathogenesis and therapy. Proc Natl Acad Sci U S A 97, 571–576. 10.1073/pnas.97.2.571.

14. Cox, D., Chang, P., Zhang, Q., Reddy, P.G., Bokoch, G.M., and Greenberg, S. (1997). Requirements for both Rac1 and Cdc42 in membrane ruffling and phagocytosis in leukocytes. J Exp Med 186, 1487–1494. 10.1084/jem.186.9.1487.

15. Danzer, K.M., Haasen, D., Karow, A.R., Moussaud, S., Habeck, M., Giese, A., Kretzschmar, H., Hengerer, B., and Kostka, M. (2007). Different species of alpha-synuclein oligomers induce calcium influx and seeding. J Neurosci 27, 9220–9232. 10.1523/JNEUROSCI.2617-07.2007.

16. Datta, A., Kim, H., Lal, M., McGee, L., Johnson, A., Moustafa, A.A., Jones, J.C., Mondal, D., Ferrer, M., and Abdel-Mageed, A.B. (2017). Manumycin A suppresses exosome biogenesis and secretion via targeted inhibition of Ras/Raf/ERK1/2 signaling and hnRNP H1 in castration-resistant prostate cancer cells. Cancer Lett 408, 73–81. 10.1016/j.canlet.2017.08.020.

17. Del Cid Pellitero, E., Shlaifer, R., Luo, W., Krahn, A., Nguyen-Renou, E., Manecka, D.-L., Rao T., Beitel, L., and Durcan, T.M. (2019). Quality Control Characterization of α-Synuclein Preformed Fibrils (PFFs). Zenodo. https://doi.org/10.5281/zenodo.3738340.

18. Desplats, P., Lee, H.-J., Bae, E.-J., Patrick, C., Rockenstein, E., Crews, L., Spencer, B., Masliah, E., and Lee, S.-J. (2009a). Inclusion formation and neuronal cell death through neuron-to-neuron transmission of α-synuclein. Proceedings of the National Academy of Sciences 106, 13010–13015. 10.1073/pnas.0903691106.

19. Desplats, P., Lee, H.J., Bae, E.J., Patrick, C., Rockenstein, E., Crews, L., Spencer, B., Masliah, E., and Lee, S.J. (2009b). Inclusion formation and neuronal cell death through neuron-to-neuron transmission of alpha-synuclein. Proc Natl Acad Sci U S A 106, 13010–13015. 10.1073/pnas.0903691106.

20. Dieriks, B.V., Park, T.I., Fourie, C., Faull, R.L., Dragunow, M., and Curtis, M.A. (2017). alpha-synuclein transfer through tunneling nanotubes occurs in SH-SY5Y cells and primary brain pericytes from Parkinson’s disease patients. Sci Rep 7, 42984. 10.1038/srep42984.

21. Erami, Z., Khalil, B.D., Salloum, G., Yao, Y., LoPiccolo, J., Shymanets, A., Nürnberg, B., Bresnick, A.R., and Backer, J.M. (2017). Rac1-stimulated macropinocytosis enhances Gβγ activation of PI3Kβ. Biochemical Journal 474, 3903–3914.

22. Fenyi, A., Leclair-Visonneau, L., Clairembault, T., Coron, E., Neunlist, M., Melki, R., Derkinderen, P., and Bousset, L. (2019). Detection of alpha-synuclein aggregates in gastrointestinal biopsies by protein misfolding cyclic amplification. Neurobiol Dis 129, 38–43. 10.1016/j.nbd.2019.05.002.

23. Fink, A.L. (2006). The Aggregation and Fibrillation of α-Synuclein. Accounts of Chemical Research 39, 628–634. 10.1021/ar050073t.

24. Fortin, D.L., Troyer, M.D., Nakamura, K., Kubo, S., Anthony, M.D., and Edwards, R.H. (2004). Lipid rafts mediate the synaptic localization of alpha-synuclein. J Neurosci 24, 6715–6723. 10.1523/JNEUROSCI.1594-04.2004.

25. Fujiwara, I., Zweifel, M.E., Courtemanche, N., and Pollard, T.D. (2018). Latrunculin A Accelerates Actin Filament Depolymerization in Addition to Sequestering Actin Monomers. Curr Biol 28, 3183–3192 e3182. 10.1016/j.cub.2018.07.082.

26. Furstner, A., De Souza, D., Turet, L., Fenster, M.D., Parra-Rapado, L., Wirtz, C., Mynott, R., and Lehmann, C.W. (2007). Total syntheses of the actin-binding macrolides latrunculin A, B, C, M, S and 16-epi-latrunculin B. Chemistry 13, 115-134. 10.1002/chem.200601135.

27. Galvez, T., Teruel, M.N., Heo, W.D., Jones, J.T., Kim, M.L., Liou, J., Myers, J.W., and Meyer, T. (2007). siRNA screen of the human signaling proteome identifies the PtdIns(3,4,5)P3-mTOR signaling pathway as a primary regulator of transferrin uptake. Genome Biol 8, R142. 10.1186/gb-2007-8-7-r142.

28. Gelpi, E., Navarro-Otano, J., Tolosa, E., Gaig, C., Compta, Y., Rey, M.J., Marti, M.J., Hernandez, I., Valldeoriola, F., Rene, R., and Ribalta, T. (2014). Multiple organ involvement by alpha-synuclein pathology in Lewy body disorders. Mov Disord 29, 1010–1018. 10.1002/mds.25776.

29. Grimmer, S., van Deurs, B., and Sandvig, K. (2002). Membrane ruffling and macropinocytosis in A431 cells require cholesterol. 115, 2953-2962.

30. Grozdanov, V., and Danzer, K.M. (2018). Release and uptake of pathologic alpha-synuclein. Cell Tissue Res 373, 175–182. 10.1007/s00441-017-2775-9.

31. Gu, C., Yaddanapudi, S., Weins, A., Osborn, T., Reiser, J., Pollak, M., Hartwig, J., and Sever, S. (2010). Direct dynamin-actin interactions regulate the actin cytoskeleton. EMBO J 29, 3593–3606. 10.1038/emboj.2010.249.

32. Hansen, C., Angot, E., Bergstrom, A.L., Steiner, J.A., Pieri, L., Paul, G., Outeiro, T.F., Melki, R., Kallunki, P., Fog, K., et al. (2011). alpha-Synuclein propagates from mouse brain to grafted dopaminergic neurons and seeds aggregation in cultured human cells. J Clin Invest 121, 715–725. 10.1172/JCI43366.

33. Holmes, B.B., DeVos, S.L., Kfoury, N., Li, M., Jacks, R., Yanamandra, K., Ouidja, M.O., Brodsky, F.M., Marasa, J., Bagchi, D.P., et al. (2013). Heparan sulfate proteoglycans mediate internalization and propagation of specific proteopathic seeds. Proc Natl Acad Sci U S A 110, E3138–3147. 10.1073/pnas.1301440110.

34. Huotari, J., and Helenius, A. (2011). Endosome maturation. EMBO J 30, 3481–3500. 10.1038/emboj.2011.286.

35. Ihse, E., Yamakado, H., van Wijk, X.M., Lawrence, R., Esko, J.D., and Masliah, E. (2017). Cellular internalization of alpha-synuclein aggregates by cell surface heparan sulfate depends on aggregate conformation and cell type. Sci Rep 7, 9008. 10.1038/s41598-017-08720-5.

36. Jao, C.C., Der-Sarkissian, A., Chen, J., and Langen, R. (2004). Structure of membrane-bound alpha-synuclein studied by site-directed spin labeling. Proc Natl Acad Sci U S A 101, 8331–8336. 10.1073/pnas.0400553101.

37. Jefri, M., Bell, S., Peng, H., Hettige, N., Maussion, G., Soubannier, V., Wu, H., Silveira, H., Theroux, J.F., Moquin, L., et al. (2020). Stimulation of L-type calcium channels increases tyrosine hydroxylase and dopamine in ventral midbrain cells induced from somatic cells. Stem Cells Transl Med 9, 697–712. 10.1002/sctm.18-0180.

38. Karpowicz, R.J., Jr., Haney, C.M., Mihaila, T.S., Sandler, R.M., Petersson, E.J., and Lee, V.M. (2017). Selective imaging of internalized proteopathic alpha-synuclein seeds in primary neurons reveals mechanistic insight into transmission of synucleinopathies. J Biol Chem 292, 13482–13497. 10.1074/jbc.M117.780296.

39. Kawahata, I., Sekimori, T., Wang, H., Wang, Y., Sasaoka, T., Bousset, L., Melki, R., Mizobata, T., Kawata, Y., and Fukunaga, K. (2021). Dopamine D2 Long Receptors Are Critical for Caveolae-Mediated alpha-Synuclein Uptake in Cultured Dopaminergic Neurons. Biomedicines 9. 10.3390/biomedicines9010049.

40. Kim, M.L., Sorg, I., and Arrieumerlou, C. (2011). Endocytosis-independent function of clathrin heavy chain in the control of basal NF-kappaB activation. PLoS One 6, e17158. 10.1371/journal.pone.0017158.

41. Koivusalo, M., Welch, C., Hayashi, H., Scott, C.C., Kim, M., Alexander, T., Touret, N., Hahn, K.M., and Grinstein, S. (2010). Amiloride inhibits macropinocytosis by lowering submembranous pH and preventing Rac1 and Cdc42 signaling. J Cell Biol 188, 547–563. 10.1083/jcb.200908086.

42. Konno, M., Hasegawa, T., Baba, T., Miura, E., Sugeno, N., Kikuchi, A., Fiesel, F.C., Sasaki, T., Aoki, M., Itoyama, Y., and Takeda, A. (2012). Suppression of dynamin GTPase decreases alpha-synuclein uptake by neuronal and oligodendroglial cells: a potent therapeutic target for synucleinopathy. Mol Neurodegener 7, 38. 10.1186/1750-1326-7-38.

43. Krueger, E.W., Orth, J.D., Cao, H., and McNiven, M.A. (2003). A dynamin-cortactin-Arp2/3 complex mediates actin reorganization in growth factor-stimulated cells. Mol Biol Cell 14, 1085–1096. 10.1091/mbc.e02-08-0466.

44. Kumar, S.T., Jagannath, S., Francois, C., Vanderstichele, H., Stoops, E., and Lashuel, H.A. (2020). How specific are the conformation-specific alpha-synuclein antibodies? Characterization and validation of 16 alpha-synuclein conformation-specific antibodies using well-characterized preparations of alpha-synuclein monomers, fibrils and oligomers with distinct structures and morphology. Neurobiol Dis 146, 105086. 10.1016/j.nbd.2020.105086.

45. Lam, H.T., Graber, M.C., Gentry, K.A., and Bieschke, J. (2016). Stabilization of α-Synuclein Fibril Clusters Prevents Fragmentation and Reduces Seeding Activity and Toxicity. Biochemistry 55, 675–685. 10.1021/acs.biochem.5b01168.

46. Lazaro, D.F., Dias, M.C., Carija, A., Navarro, S., Madaleno, C.S., Tenreiro, S., Ventura, S., and Outeiro, T.F. (2016). The effects of the novel A53E alpha-synuclein mutation on its oligomerization and aggregation. Acta Neuropathol Commun 4, 128. 10.1186/s40478-016-0402-8.

47. Lee, H.J., Choi, C., and Lee, S.J. (2002). Membrane-bound alpha-synuclein has a high aggregation propensity and the ability to seed the aggregation of the cytosolic form. J Biol Chem 277, 671–678. 10.1074/jbc.M107045200.

48. Lee, H.J., Suk, J.E., Bae, E.J., Lee, J.H., Paik, S.R., and Lee, S.J. (2008). Assembly-dependent endocytosis and clearance of extracellular alpha-synuclein. Int J Biochem Cell Biol 40, 1835–1849. 10.1016/j.biocel.2008.01.017.

49. Lee, H.J., Suk, J.E., Patrick, C., Bae, E.J., Cho, J.H., Rho, S., Hwang, D., Masliah, E., and Lee, S.J. (2010). Direct transfer of alpha-synuclein from neuron to astroglia causes inflammatory responses in synucleinopathies. J Biol Chem 285, 9262–9272. 10.1074/jbc.M109.081125.

50. Lewy, F. (1912). Handbuch der Neurologie (Julius Springer).

51. Li, J., Uversky, V.N., and Fink, A.L. (2002). Conformational Behavior of Human α-Synuclein is Modulated by Familial Parkinson’s Disease Point Mutations A30P and A53T. NeuroToxicology 23, 553–567. 10.1016/s0161-813x(02)00066-9.

52. Li, J.Y., Englund, E., Holton, J.L., Soulet, D., Hagell, P., Lees, A.J., Lashley, T., Quinn, N.P., Rehncrona, S., Bjorklund, A., et al. (2008). Lewy bodies in grafted neurons in subjects with Parkinson’s disease suggest host-to-graft disease propagation. Nat Med 14, 501–503. 10.1038/nm1746.

53. Li, J.Y., Englund, E., Widner, H., Rehncrona, S., Bjorklund, A., Lindvall, O., and Brundin, P. (2010). Characterization of Lewy body pathology in 12- and 16-year-old intrastriatal mesencephalic grafts surviving in a patient with Parkinson’s disease. Mov Disord 25, 1091–1096. 10.1002/mds.23012.

54. Li, L., Wan, T., Wan, M., Liu, B., Cheng, R., and Zhang, R. (2015). The effect of the size of fluorescent dextran on its endocytic pathway. Cell Biol Int 39, 531–539. 10.1002/cbin.10424.

55. Lim, C.Y., Davis, O.B., Shin, H.R., Zhang, J., Berdan, C.A., Jiang, X., Counihan, J.L., Ory, D.S., Nomura, D.K., and Zoncu, R. (2019). ER-lysosome contacts enable cholesterol sensing by mTORC1 and drive aberrant growth signalling in Niemann-Pick type C. Nat Cell Biol 21, 1206-1218. 10.1038/s41556-019-0391-5.

56. Liu, J., Zhou, Y., Wang, Y., Fong, H., Murray, T.M., and Zhang, J. (2007). Identification of proteins involved in microglial endocytosis of alpha-synuclein. J Proteome Res 6, 3614–3627. 10.1021/pr0701512.

57. Luk, K.C., Covell, D.J., Kehm, V.M., Zhang, B., Song, I.Y., Byrne, M.D., Pitkin, R.M., Decker, S.C., Trojanowski, J.Q., and Lee, V.M. (2016). Molecular and Biological Compatibility with Host Alpha-Synuclein Influences Fibril Pathogenicity. Cell Rep 16, 3373–3387. 10.1016/j.celrep.2016.08.053.

58. Luk, K.C., Kehm, V., Carroll, J., Zhang, B., O’Brien, P., Trojanowski, J.Q., and Lee, V.M. (2012). Pathological alpha-synuclein transmission initiates Parkinson-like neurodegeneration in nontransgenic mice. Science 338, 949–953. 10.1126/science.1227157.

59. Luk, K.C., Song, C., O’Brien, P., Stieber, A., Branch, J.R., Brunden, K.R., Trojanowski, J.Q., and Lee, V.M. (2009). Exogenous alpha-synuclein fibrils seed the formation of Lewy body-like intracellular inclusions in cultured cells. Proc Natl Acad Sci U S A 106, 20051–20056. 10.1073/pnas.0908005106.

60. Madeira, A., Yang, J., Zhang, X., Vikeved, E., Nilsson, A., Andren, P.E., and Svenningsson, P. (2011). Caveolin-1 interacts with alpha-synuclein and mediates toxic actions of cellular alpha-synuclein overexpression. Neurochem Int 59, 280–289. 10.1016/j.neuint.2011.05.017.

61. Maneca, D.-L., Luo, W., Krahn, A., Del Cid Pellitero, E., Shlaifer, I., Beitel, L.K., Rao, T., and Durcan, T.M. (2019). Production of Recombinant α Synuclein Monomers and Preformed Fibrils (PFFs). Zenodo. http://doi.org/10.5281/zenodo.3738335.

62. Manning-Bog, A.B., McCormack, A.L., Li, J., Uversky, V.N., Fink, A.L., and Di Monte, D.A. (2002). The herbicide paraquat causes up-regulation and aggregation of alpha-synuclein in mice: paraquat and alpha-synuclein. J Biol Chem 277, 1641–1644. 10.1074/jbc.C100560200.

63. Mao, X., Ou, M.T., Karuppagounder, S.S., Kam, T.I., Yin, X., Xiong, Y., Ge, P., Umanah, G.E., Brahmachari, S., Shin, J.H., et al. (2016). Pathological alpha-synuclein transmission initiated by binding lymphocyte-activation gene 3. Science 353. 10.1126/science.aah3374.

64. Masaracchia, C., Hnida, M., Gerhardt, E., Lopes da Fonseca, T., Villar-Pique, A., Branco, T., Stahlberg, M.A., Dean, C., Fernandez, C.O., Milosevic, I., and Outeiro, T.F. (2018). Membrane binding, internalization, and sorting of alpha-synuclein in the cell. Acta Neuropathol Commun 6, 79. 10.1186/s40478-018-0578-1.

65. Masuda-Suzukake, M., Nonaka, T., Hosokawa, M., Kubo, M., Shimozawa, A., Akiyama, H., and Hasegawa, M. (2014). Pathological alpha-synuclein propagates through neural networks. Acta Neuropathol Commun 2, 88. 10.1186/s40478-014-0088-8.

66. Masuda-Suzukake, M., Nonaka, T., Hosokawa, M., Oikawa, T., Arai, T., Akiyama, H., Mann, D.M., and Hasegawa, M. (2013). Prion-like spreading of pathological alpha-synuclein in brain. Brain 136, 1128–1138. 10.1093/brain/awt037.

67. Mayor, S., and Pagano, R.E. (2007). Pathways of clathrin-independent endocytosis. Nat Rev Mol Cell Biol 8, 603–612. 10.1038/nrm2216.

68. Mills, I.G., Jones, A.T., and Clague, M.J. (1998). Involvement of the endosomal autoantigen EEA1 in homotypic fusion of early endosomes. Current Biology 8, 881–884. 10.1016/s0960-9822(07)00351-x.

69. Morton, W.M., Ayscough, K.R., and McLaughlin, P.J. (2000). Latrunculin alters the actin-monomer subunit interface to prevent polymerization. Nat Cell Biol 2, 376–378. 10.1038/35014075.

70. Mougenot, A.L., Nicot, S., Bencsik, A., Morignat, E., Verchere, J., Lakhdar, L., Legastelois, S., and Baron, T. (2012). Prion-like acceleration of a synucleinopathy in a transgenic mouse model. Neurobiol Aging 33, 2225–2228. 10.1016/j.neurobiolaging.2011.06.022.

71. Mulherkar, N., Raaben, M., de la Torre, J.C., Whelan, S.P., and Chandran, K. (2011). The Ebola virus glycoprotein mediates entry via a non-classical dynamin-dependent macropinocytic pathway. Virology 419, 72–83. 10.1016/j.virol.2011.08.009.

72. Nakamura, K., Mori, F., Kon, T., Tanji, K., Miki, Y., Tomiyama, M., Kurotaki, H., Toyoshima, Y., Kakita, A., Takahashi, H., et al. (2015). Filamentous aggregations of phosphorylated alpha-synuclein in Schwann cells (Schwann cell cytoplasmic inclusions) in multiple system atrophy. Acta Neuropathol Commun 3, 29. 10.1186/s40478-015-0208-0.

73. Nakamura, K., Nemani, V.M., Wallender, E.K., Kaehlcke, K., Ott, M., and Edwards, R.H. (2008). Optical reporters for the conformation of alpha-synuclein reveal a specific interaction with mitochondria. J Neurosci 28, 12305–12317. 10.1523/JNEUROSCI.3088-08.2008.

74. Nakase, I., Kobayashi, N.B., Takatani-Nakase, T., and Yoshida, T. (2015). Active macropinocytosis induction by stimulation of epidermal growth factor receptor and oncogenic Ras expression potentiates cellular uptake efficacy of exosomes. Sci Rep 5, 10300. 10.1038/srep10300.

75. Oh, P., McIntosh, D.P., and Schnitzer, J.E. (1998). Dynamin at the neck of caveolae mediates their budding to form transport vesicles by GTP-driven fission from the plasma membrane of endothelium. J Cell Biol 141, 101–114. 10.1083/jcb.141.1.101.

76. Oh, S.H., Kim, H.N., Park, H.J., Shin, J.Y., Bae, E.J., Sunwoo, M.K., Lee, S.J., and Lee, P.H. (2016). Mesenchymal Stem Cells Inhibit Transmission of alpha-Synuclein by Modulating Clathrin-Mediated Endocytosis in a Parkinsonian Model. Cell Rep 14, 835–849. 10.1016/j.celrep.2015.12.075.

77. Park, J.Y., Kim, K.S., Lee, S.B., Ryu, J.S., Chung, K.C., Choo, Y.K., Jou, I., Kim, J., and Park, S.M. (2009). On the mechanism of internalization of alpha-synuclein into microglia: roles of ganglioside GM1 and lipid raft. J Neurochem 110, 400–411. 10.1111/j.1471-4159.2009.06150.x.

78. Park, R.J., Shen, H., Liu, L., Liu, X., Ferguson, S.M., and De Camilli, P. (2013). Dynamin triple knockout cells reveal off target effects of commonly used dynamin inhibitors. J Cell Sci 126, 5305–5312. 10.1242/jcs.138578.

79. Pelkmans, L., Puntener, D., and Helenius, A. (2002). Local actin polymerization and dynamin recruitment in SV40-induced internalization of caveolae. Science 296, 535–539. 10.1126/science.1069784.

80. Pieri, L., Madiona, K., and Melki, R. (2016). Structural and functional properties of prefibrillar alpha-synuclein oligomers. Sci Rep 6, 24526. 10.1038/srep24526.

81. Preta, G., Cronin, J.G., and Sheldon, I.M. (2015). Dynasore - not just a dynamin inhibitor. Cell Commun Signal 13, 24. 10.1186/s12964-015-0102-1.

82. Recasens, A., and Dehay, B. (2014). Alpha-synuclein spreading in Parkinson’s disease. Front Neuroanat 8, 159. 10.3389/fnana.2014.00159.

83. Recasens, A., Ulusoy, A., Kahle, P.J., Di Monte, D.A., and Dehay, B. (2018). In vivo models of alpha-synuclein transmission and propagation. Cell Tissue Res 373, 183–193. 10.1007/s00441-017-2730-9.

84. Reyes, J.F., Rey, N.L., Bousset, L., Melki, R., Brundin, P., and Angot, E. (2014). Alpha-synuclein transfers from neurons to oligodendrocytes. Glia 62, 387–398. 10.1002/glia.22611.

85. Rodriguez, L., Marano, M.M., and Tandon, A. (2018). Import and Export of Misfolded alpha-Synuclein. Front Neurosci 12, 344. 10.3389/fnins.2018.00344.

86. Rutherford, N.J., Moore, B.D., Golde, T.E., and Giasson, B.I. (2014). Divergent effects of the H50Q and G51D SNCA mutations on the aggregation of alpha-synuclein. J Neurochem 131, 859–867. 10.1111/jnc.12806.

87. Samuel, F., Flavin, W.P., Iqbal, S., Pacelli, C., Sri Renganathan, S.D., Trudeau, L.E., Campbell, E.M., Fraser, P.E., and Tandon, A. (2016). Effects of Serine 129 Phosphorylation on alpha-Synuclein Aggregation, Membrane Association, and Internalization. J Biol Chem 291, 4374–4385. 10.1074/jbc.M115.705095.

88. Sandal, M., Valle, F., Tessari, I., Mammi, S., Bergantino, E., Musiani, F., Brucale, M., Bubacco, L., and Samori, B. (2008). Conformational equilibria in monomeric alpha-synuclein at the single-molecule level. PLoS Biol 6, e6. 10.1371/journal.pbio.0060006.

89. Schneider, C.A., Rasband, W.S., and Eliceiri, K.W. (2012). NIH Image to ImageJ: 25 years of image analysis. Nat Methods 9, 671–675. 10.1038/nmeth.2089.

90. Shahmoradian, S.H., Lewis, A.J., Genoud, C., Hench, J., Moors, T.E., Navarro, P.P., Castano-Diez, D., Schweighauser, G., Graff-Meyer, A., Goldie, K.N., et al. (2019). Lewy pathology in Parkinson’s disease consists of crowded organelles and lipid membranes. Nat Neurosci 22, 1099–1109. 10.1038/s41593-019-0423-2.

91. Shearer, L.J., Petersen, N.O., and Woodside, M.T. (2021). Internalization of alpha-synuclein oligomers into SH-SY5Y cells. Biophys J. 10.1016/j.bpj.2020.12.031.

92. Sigismund, S., Woelk, T., Puri, C., Maspero, E., Tacchetti, C., Transidico, P., Di Fiore, P.P., and Polo, S. (2005). Clathrin-independent endocytosis of ubiquitinated cargos. Proc Natl Acad Sci U S A 102, 2760–2765. 10.1073/pnas.0409817102.

93. Spillantini, M.G., Schmidt, M.L., Lee, V.M.-Y., Trojanowski, J.Q., Jakes, R., and Goedert, M.J.N. (1997). α-Synuclein in Lewy bodies. 388, 839–840.

94. Sung, J.Y., Kim, J., Paik, S.R., Park, J.H., Ahn, Y.S., and Chung, K.C. (2001). Induction of neuronal cell death by Rab5A-dependent endocytosis of alpha-synuclein. J Biol Chem 276, 27441–27448. 10.1074/jbc.M101318200.

95. Tanaka, M., Kim, Y.M., Lee, G., Junn, E., Iwatsubo, T., and Mouradian, M.M. (2004). Aggresomes formed by alpha-synuclein and synphilin-1 are cytoprotective. J Biol Chem 279, 4625–4631. 10.1074/jbc.M310994200.

96. Trischler, M., Stoorvogel, W., and Ullrich, O. (1999). Biochemical analysis of distinct Rab5- and Rab11-positive endosomes along the transferrin pathway. 112, 4773-4783.

97. Uemura, N., Uemura, M.T., Luk, K.C., Lee, V.M., and Trojanowski, J.Q. (2020). Cell-to-Cell Transmission of Tau and alpha-Synuclein. Trends Mol Med. 10.1016/j.molmed.2020.03.012.

98. Uemura, N., Yagi, H., Uemura, M.T., Hatanaka, Y., Yamakado, H., and Takahashi, R. (2018). Inoculation of alpha-synuclein preformed fibrils into the mouse gastrointestinal tract induces Lewy body-like aggregates in the brainstem via the vagus nerve. Mol Neurodegener 13, 21. 10.1186/s13024-018-0257-5.

99. Underwood, R., Wang, B., Pathak, A., Volpicelli-Daley, L., and Yacoubian, T.A. (2020). Rab27 GTPases regulate alpha-synuclein uptake, cell-to-cell transmission, and toxicity. bioRxiv. 10.1101/2020.11.17.387449.

100. Uversky, V.N. (2007). Neuropathology, biochemistry, and biophysics of alpha-synuclein aggregation. J Neurochem 103, 17–37. 10.1111/j.1471-4159.2007.04764.x.

101. Vargas, K.J., Schrod, N., Davis, T., Fernandez-Busnadiego, R., Taguchi, Y.V., Laugks, U., Lucic, V., and Chandra, S.S. (2017). Synucleins Have Multiple Effects on Presynaptic Architecture. Cell Rep 18, 161–173. 10.1016/j.celrep.2016.12.023.

102. Volpicelli-Daley, L.A., Luk, K.C., and Lee, V.M. (2014). Addition of exogenous alpha-synuclein preformed fibrils to primary neuronal cultures to seed recruitment of endogenous alpha-synuclein to Lewy body and Lewy neurite-like aggregates. Nat Protoc 9, 2135–2146. 10.1038/nprot.2014.143.

103. Volpicelli-Daley, L.A., Luk, K.C., Patel, T.P., Tanik, S.A., Riddle, D.M., Stieber, A., Meaney, D.F., Trojanowski, J.Q., and Lee, V.M. (2011). Exogenous alpha-synuclein fibrils induce Lewy body pathology leading to synaptic dysfunction and neuron death. Neuron 72, 57–71. 10.1016/j.neuron.2011.08.033.

104. Wakatsuki, T., Schwab, B., Thompson, N.C., and Elson, E.L. (2001). Effects of cytochalasin D and latrunculin B on mechanical properties of cells. Journal of Cell Science 114, 1025–1036.

105. Westphal, C.H., and Chandra, S.S. (2013). Monomeric synucleins generate membrane curvature. J Biol Chem 288, 1829–1840. 10.1074/jbc.M112.418871.

106. Williams, T.D., and Kay, R.R. (2018). The physiological regulation of macropinocytosis during Dictyostelium growth and development. 131, jcs213736. 10.1242/jcs.213736 %J Journal of Cell Science.

107. Yan, F., Chen, Y., Li, M., Wang, Y., Zhang, W., Chen, X., and Ye, Q. (2018). Gastrointestinal nervous system alpha-synuclein as a potential biomarker of Parkinson disease. Medicine (Baltimore) 97, e11337. 10.1097/MD.0000000000011337.

108. Yoshida, S., Pacitto, R., Inoki, K., and Swanson, J. (2018). Macropinocytosis, mTORC1 and cellular growth control. Cell Mol Life Sci 75, 1227-1239. 10.1007/s00018-017-2710-y.

109. Zeineddine, R., Pundavela, J.F., Corcoran, L., Stewart, E.M., Do-Ha, D., Bax, M., Guillemin, G., Vine, K.L., Hatters, D.M., Ecroyd, H., et al. (2015). SOD1 protein aggregates stimulate macropinocytosis in neurons to facilitate their propagation. Mol Neurodegener 10, 57. 10.1186/s13024-015-0053-4.

110. Zhang, Q., Xu, Y., Lee, J., Jarnik, M., Wu, X., Bonifacino, J.S., Shen, J., and Ye, Y. (2020). A myosin-7B-dependent endocytosis pathway mediates cellular entry of alpha-synuclein fibrils and polycation-bearing cargos. Proc Natl Acad Sci U S A 117, 10865–10875. 10.1073/pnas.1918617117.

111. Zhang, W., Wang, T., Pei, Z., Miller, D.S., Wu, X., Block, M.L., Wilson, B., Zhang, W., Zhou, Y., Hong, J.S., and Zhang, J. (2005). Aggregated alpha-synuclein activates microglia: a process leading to disease progression in Parkinson’s disease. FASEB J 19, 533–542. 10.1096/fj.04-2751com.

112. Zwartkruis, F.J., and Burgering, B.M. (2013). Ras and macropinocytosis: trick and treat. Cell Res 23, 982–983. 10.1038/cr.2013.79.

