## Supplemental Materials for "Rapid macropinocytic transfer of α-synuclein to lysosomes"

### SUPPLEMENTARY MATERIALS

#### Table S1: Previous studies on α-syn endocytosis

| Type of α-syn | Inhibition/Inhibitors used | Incubation time following α-syn addition | Reported Intracellular trafficking | Findings | Publication |
|---|---|---|---|---|---|
| Monomer | | 1 h | Rab5A | Rab5A-mediated uptake of α-syn | (Sung *et al.*, 2001) |
| Fibril | Cytochalasin D | 2 and 12 h | | Cytochalasin D treatment resulted in decreased α-syn uptake in microglia | (Zhang et al., 2005) |
| Fibril | | 3 h | | Co-immunoprecipitation of clathrin and α-syn; colocalization of α-syn and clathrin | (Liu *et al.*, 2007) |
| Fibril | dynamin-1 K44A | 1 h | Partial colocalization of α-syn with EEA1 and LAMP2. | Internalization of α-syn fibrils was inhibited by low temperature and dynamin dominant-negative, while monomeric α-syn entered cells through diffusion. It was concluded that the endolysosomal pathway is involved. | (Lee *et al.*, 2008) |
| Monomer | Dynamin dominant negative and dynasore | 1 h | | Internalization of α-syn was not inhibited with dynamin dominant-negative or dynasore but lipid-raft mediated. | (Park et al., 2009) |
| Cellular synuclein expression | Dynamin-1 K44A (dominant negative) | 24 and 48 h | | Synuclein transmission was inhibited with the use of dynamin dominant-negative | (Desplats *et al.*, 2009b) |
| Vector expression of α-syn | Dynamin dominant-negative | Coculture – 3 days | Endolysosomal system | α-syn endocytosis is inhibited in cells expressing dynamin dominant-negative | (Lee et al., 2010) |
| Transfected and exogenous α-syn monomer | Dynasore, monodansylcadaverine | Coculture, injection in rat cortex | | α-syn endocytosis and transfer occur in a dynamin-dependent manner | (Hansen *et al.*, 2011) |
| Monomeric α-syn | siRNA knockdown of dynamin, inhibition of dynamin GTPase activity by sertraline | 24 h | Rab5A and LAMP1 | Synuclein endocytosis was decreased by inhibiting dynamin GTPases | (Konno *et al.*, 2012) |
| Tau and α-syn fibrils, TAT | Inhibition of tau uptake via Cytochalasin, Latrunculin, Rottlerin, Amiloride. Tau uptake was not inhibited by dynasore. α-syn uptake was inhibited by heparin and chlorate | 3 and 5 h | | When administered together, tau, TAT, and α-syn colocalize, suggesting α-syn fibrils are endocytosed via macropinocytosis. Tau fibrils did not colocalize with the clathrin antibody. | (Holmes et al., 2013) |
| Monomer, oligomer, fibril | Dynasore | 0,3,6,12,24 h | | Dynasore inhibited synuclein uptake in a concentration-dependent manner | (Reyes et al., 2014) |
| Transfection: α-syn expression | Dynasore, Pitstop 2, | | EEA1, NR1, NR2A | Clathrin-mediated internalization of α-syn | (Oh et al., 2016) |
| Fibril | EIPA | | | α-syn induced membrane ruffling, but EIPA, a macropinocytic inhibitor, did not inhibit its internalization | (Zeineddine *et al.*, 2015) |
| Fibrils | Dynamin 1 dominant negative | 1 h | LAMP1 | Dynamin is partially required for the uptake, but not the intercellular transfer of α-syn fibrils. | (Abounit et al., 2016) |
| Fibrils and phospho-fibrils | Dynasore | 20 min | | Inhibition of clathrin-mediated endocytosis through dynasore; decreased uptake of fibrils | (Samuel et al., 2016) |
| Fibrils | LAG3 KO, LAG3 antibodies | 5, 10, 20 min and more | Rab5, Rab7, LAMP1 | LAG3 mediated endocytosis of α-syn PFF, which then undergoes receptor-mediated endocytosis | (Mao *et al.*, 2016) |
| Fibrils | chloroquine | 3,15,60 min and longer | Endolysosomal pathway, LAMP1 staining | PFF is transported through the endolysosomal pathway | (Karpowicz *et al.*, 2017) |
| Fibrils | | 4 and 24 h | | Amyloid fibrils depend on Heparan sulfate for internalization | (Ihse et al., 2017) |
| Monomer and fibril | Dyngo, Pitstop | 24 h | Endosomal maturation, and finally, lysosomes | α-syn monomers use dynamin-mediated endocytosis, colocalize with Rab4, 5, and 7, and trafficking through the endosomal system. α-syn fibrils undertake a different pathway | (Masaracchia et al., 2018) |
| Monomer, oligomer and fibril | | 24 h | Colocalization with endolysosomal markers | Clathrin-mediated endocytosis of α-syn colocalization of synuclein and transferrin | (Hoffmann et al., 2019) |
| Monomer and fibril | Dynasore, caveolin-1 siRNA, | 48 h | | Caveolae-dependent uptake of α-syn in dopaminergic neurons | (Kawahata et al., 2021) |
| Various oligomers (varying sizes) | Pitstop | 2 h | Rab5 and LAMP-1 | Larger oligomers are more dependent on clathrin-mediated endocytosis, colocalization with early endosomes and lysosomes | (Shearer et al., 2021) |

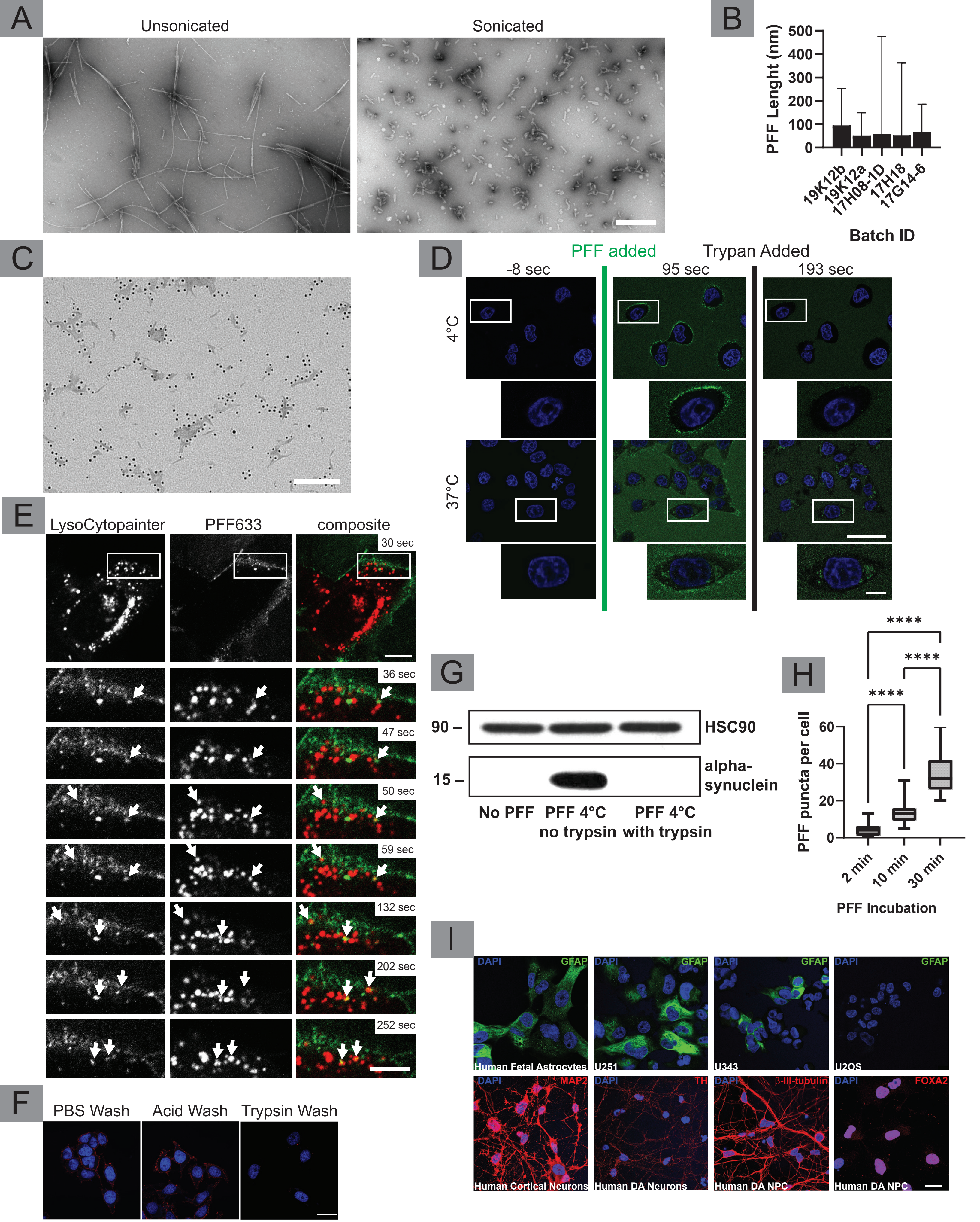


Bayati et al., Fig. S1

**Figure S1, related to Figure 1. Characterization of PFF, trypan blue exclusion assay, live PFF uptake, trypsin wash, and characterization of cell types.** (**A**) Sonication of α-syn fibrils results in significantly smaller fibril length, allowing the α-syn oligomers to be characterized as PFF with a length below 100 nm. (**B**) The mean length of unconjugated PFF samples used; mean ± upper and lower limits. Once confirmed with electron microscopy, the corresponding batches were tagged with Alexa Fluor 488, Alexa Fluor 633, or 5 nm gold (**C**). Gold-tagged PFF. Scale Bar = 100 nm. (**D**) Trypan blue exclusion assay was used to discriminate against intra- and extracellular PFF**.** HeLa cells were grown on glass-bottomed plates and placed on ice for 30 min. They were then transferred to an imaging chamber for live imaging. For the 4°C condition, the chamber was not heated: cells remained at low temperature (retrieved from ice immediately before imaging). Images were taken before and after PFF addition. Trypan blue was added to cells 2 min following PFF addition. No PFF was internalized in this condition, as trypan blue was able to quench all PFF fluorescence. For the 37°C condition, the chamber was heated to 37°C before the addition of PFF. PFF was then added, and trypan blue was added 2 min following PFF addition. PFF fluorescence not quenched by trypan blue signifies internalized PFF. Scale bar = 20 μm for low magnification and 5 μm for insets. (**E**) PFF was added to cells at 4°C for 30 min to allow PFF to bind to the cell surface without internalization. Cells were then transferred to a pre-heated live imaging chamber. Arrows point to specific PFF (green) punctae that colocalize with lysosomes (red) over time. Scale bar = 10 μm for the low magnification image and 5 μm for the insets. (**F**) Unlike cargo proteins like Tf and EGF, trypsinization for 90 sec on ice is needed to remove extracellular PFF. In this experiment, cells were incubated with PFF for 30 min at 4°C. Since PFF internalization is temperature-dependent, we expected no PFF internalization. We then assessed the efficacy of PBS, Acid, and Trypsin wash on their ability to remove extracellular PFF. Scale bar = 20 μm. (**G**) To further confirm the efficacy of trypsin wash for the removal of extracellular PFF, PBS (control) or PFF were added to cells. Cells were incubated at 4°C for 30 min. Cells exposed to PFF were then washed with either PBS or with trypsin. Cell lysates were then immunoblotted for α-syn to assess the presence of PFF. Samples washed with trypsin showed no α-syn signal, while the cells washed with PBS did, confirming the efficacy of trypsin in removing extracellular PFF. (**H**) To better visualize the buildup of PFF in cells, we also counted the number of puncta of PFF appearing in cells at different time points. n = 50 (individual cells) for each time point (n = 150 total), counted from 9 images per time point captured from three different independent experiments. Data were statistically analyzed using one-way ANOVA and Tukey for mean comparisons; the plot shows mean ± min & max values for each condition; p<0.0001 denoted as ****. (**I**) To confirm the identity of astrocytes, glioblastoma cell lines, and the iPSC-derived neural progenitor cells along with differentiated neurons, marker antibodies were used. Astrocytic and glial identity was confirmed using the glial fibrillary associated protein (GFAP) antibody. Neuronal markers such as microtubule-associated protein 2 (MAP2) were used to confirm neuronal identity. To confirm the identity of dopaminergic neurons, tyrosine hydroxylase antibody (TH) was used. The identity of neural progenitor cells was confirmed with beta-III tubulin was used. Finally, to confirm the identity of dopaminergic neural progenitor cells, Forkhead box protein A2 (FOXA2) was used. Scale bar = 20 μm.

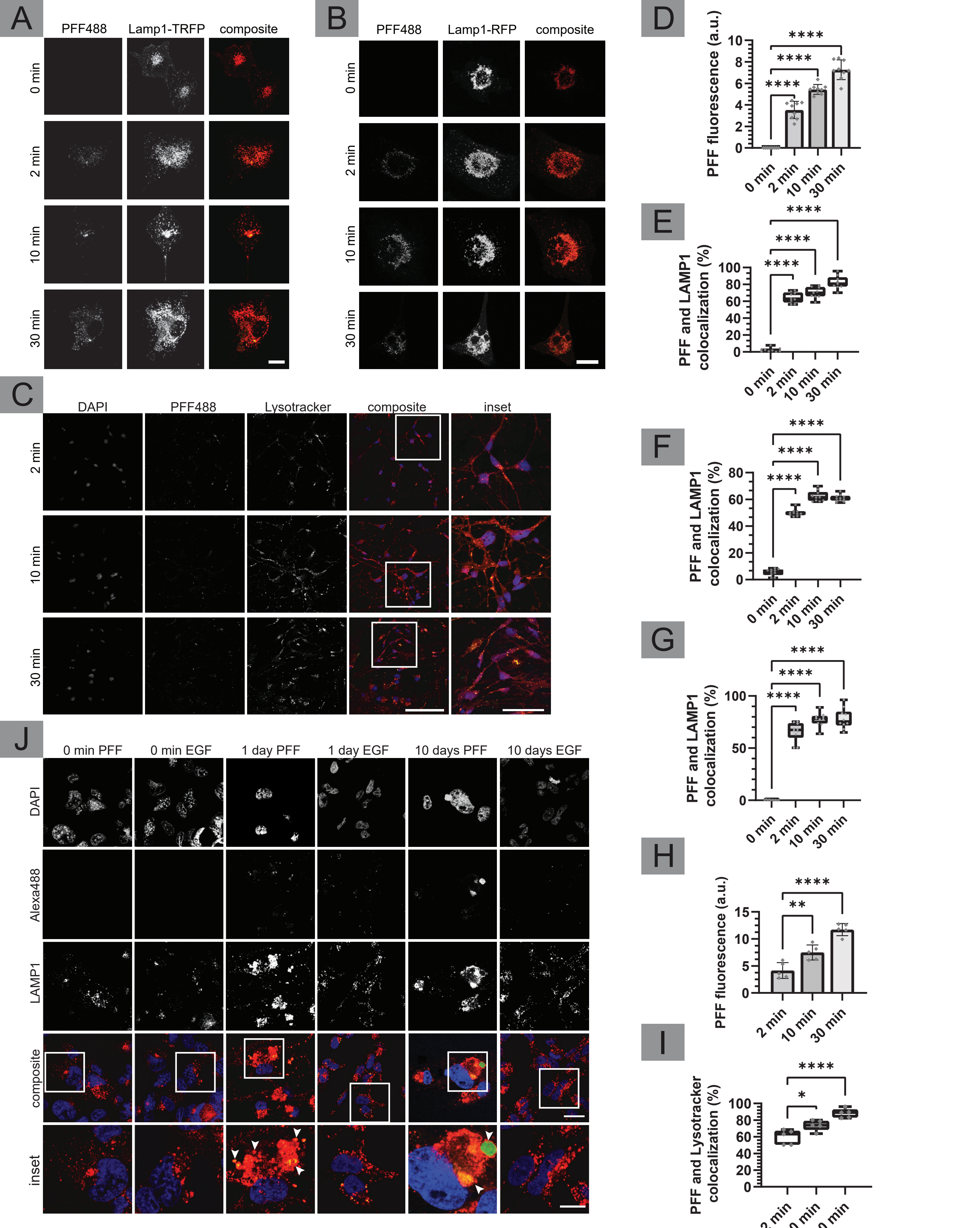


Bayati et al., Fig. S2

**Figure S2, related to Figure 2. PFF colocalization with lysosomes in glioblastomas and cortical neurons.**

(**A**) U87 cells were plated on coverslips and transfected with LAMP1-TurboRFP. PFF was added to each coverslip at 2 µg/ml. Cells were incubated for 0, 2, 10, and 30 min at 37°C following the addition of PFF. Cells were washed with trypsin and fixed. PFF (green) colocalization with LAMP1 (red) can be seen as early as 2 min. Scale bar = 20 µm. (**B**) U251 cells transduced with LAMP1-RFP lentivirus were mounted onto coverslips and were incubated with PFF as in **A**. Cells were then trypsin washed and fixed. Scale bar = 20 µm. (**C**) Cortical NPCs were mounted on coverslips. Cells were then stained with lysotracker for 30 min at 37°C. PFF was added to each coverslip at 2 µg/ml. Cells were incubated for 0, 2, 10, 30 min at 37°C following the addition of PFF. Cells were washed with trypsin and fixed. Scale bar = 20 µm, and insets are 10 µm. (**D**) Quantification of PFF uptake from experiment in **A.** n = 9 for each condition (n = 36 total); mean ± SD (**E**) Colocalization of PFF and LAMP1 in **A**. n = 9 for each condition (n = 36 total); mean ± min & max values. (**F**) Colocalization of LAMP1 with PFF, respectively, from experiment in **B**. n = 10 for each condition (n = 40 total); mean ± min & max.(**G**) Colocalization of LAMP1 with PFF from experiments done in U343 cells. n = 10 for each condition (n = 40 total); mean ± min & max. **(H)** Quantification of PFF uptake from experiment in **C.** n = 5 for each condition (n = 15); mean ± SD. (**I**) Colocalization of PFF with Lysotracker, n = 5 for each condition (n = 15 total); mean ± min & max. For all quantifications, data was collected from three independent experiments. Statistical analysis was done with one-way ANOVA and Tukey for mean comparisons; individual data points were shown with gray diamonds. $p < 0.0001$ denoted as ****, $p < 0.001$ denoted as ***, $p < 0.05$ denoted as *. (**J**) U343 human glioblastoma cells were mounted on coverslips given PFF488 or EGF488 at 2 µg/ml and 0.2 ug/ml concentration respectively on ice for 1 h. Cells were removed from the ice, given fresh media, and then incubated for 0 h, 1 day, or 10 days at 37°C. Following incubation, cells were trypsin washed and fixed. Samples were then permeabilized, blocked, and stained with LAMP1 antibody. PFF fluorescence builds over 10 days while EGF fluorescence dissipates. Arrowheads point to large PFF and LAMP1 structures, which only present in cells exposed to PFF. Scale bar = 20 µm for low magnification and 5 µm for insets.

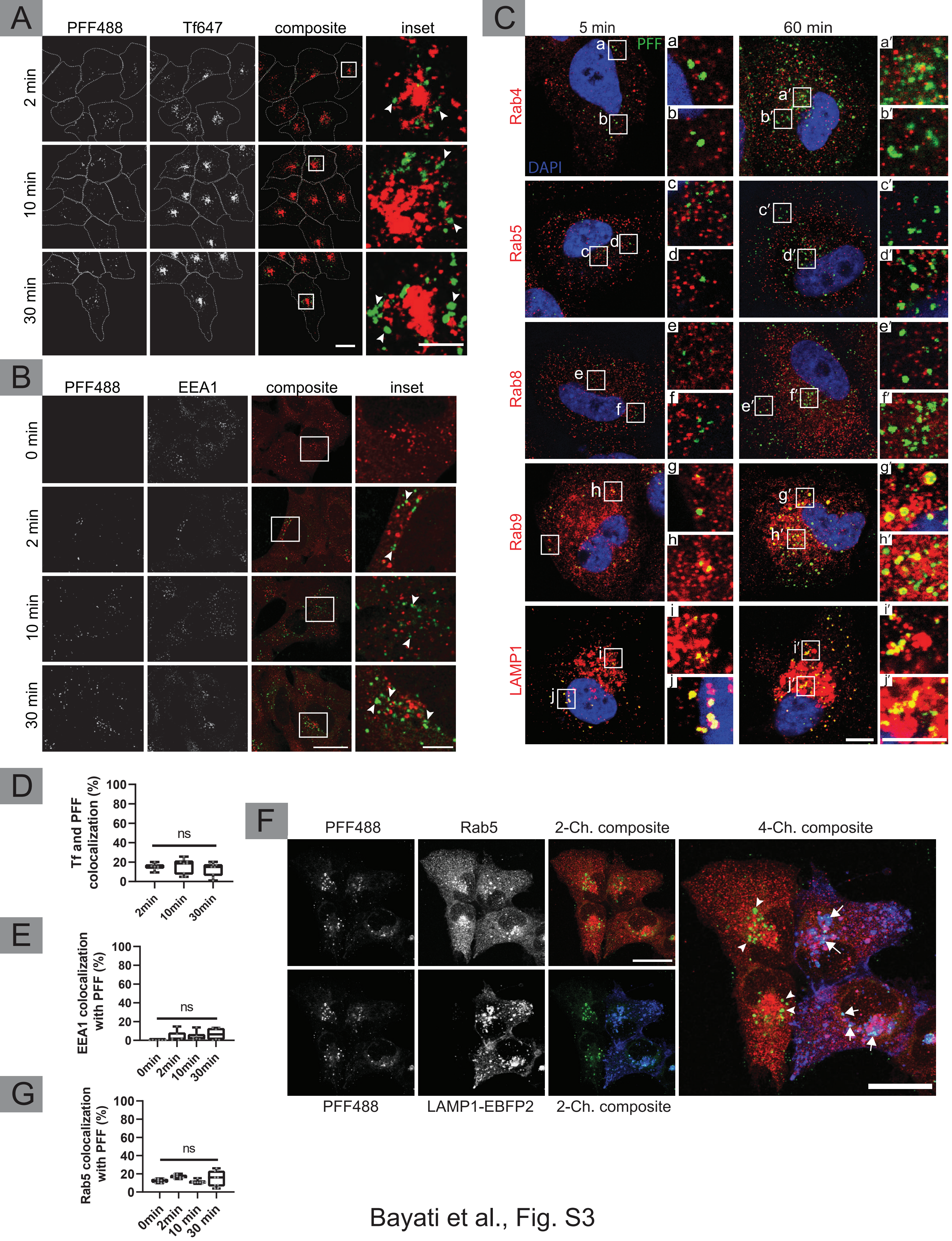


Bayati et al., Fig. S3

**Figure S3, related to Figure 3. PFF is not trafficked through early/recycling endosomes.** (**A**) PFF (green) and Tf (red) were added to U2OS cells at 2 µg/ml and 0.2 µg/ml, respectively, and the cells were incubated for 2, 10, and 30 min at 37°C. Cells were washed with trypsin and fixed. Arrowheads point to large PFF punctae. Scale bar = 20 µm and 10 µm for insets. (**B**) U2OS cells were mounted on coverslips, PFF (green) was added to each coverslip at 2 µg/ml and the cells were incubated for 0, 2, 10, and 30 min at 37°C. Cells were washed with trypsin, fixed, permeabilized, and stained with EEA1 antibody (red). Arrowheads point to large PFF punctae. Scale bar = 20 µm and 5 µm. (**C**) PFF (green) colocalization with early endosome markers: Rab4 and Rab5; Rab8; and lysosomal markers: Rab9 and LAMP1 was examined using antibodies (red). PFF colocalized only with Rab9 and LAMP1 at 5 and 60 min. The scale bar for low magnification images is 10 µm and 5 µm for insets. (**D**) Colocalization rate of Tf with PFF from experiments as in **A**. n = 9 for each condition (n = 36 total), from three independent experiments, mean ± min & max values; one-way ANOVA; $p > 0.05$ denoted as ns for not significant. (**E**) Colocalization of EEA1 with PFF from experiments in **B**. n = 6 for each condition (n = 24 total), from three independent experiments, mean ± min & max values; one-way ANOVA; $p > 0.05$ denoted as ns for not significant. (**F**) U2OS cells were incubated with PFF at 2 µg/ml for 5 minutes. Cells were washed with trypsin, fixed, permeabilized, and stained with Rab5 antibody. Scale bar = 20 µm. (**G**) Colocalization rate of Rab5 with PFF from experiments as in **F.** n = 9 for each condition (n = 36 total), from three independent experiments, mean ± min & max values; one-way ANOVA; $p > 0.05$ denoted as ns for not significant. All data points are shown using gray diamonds.

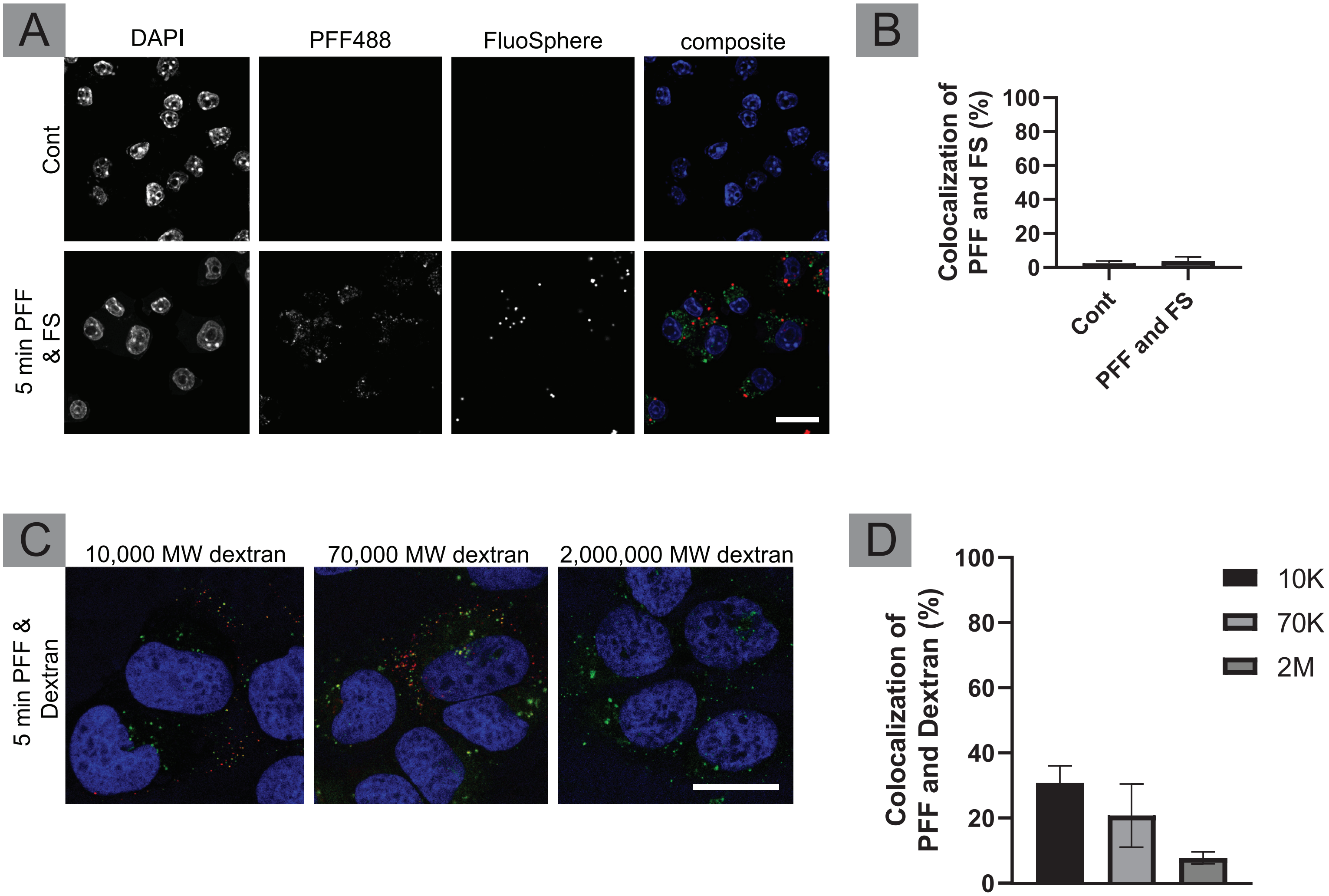


Bayati et al., Fig. S4

**Figure S4, related to Figure 4. PFF internalization follows a unique pathway different from conventional bulk and fluid-phase endocytosis.**
(**A**) RAW 264.7 cells, monocyte/macrophage-like cells, were plated on coverslips and administered FluoSpheres (FS) with 1 um diameter alongside PFF. Scale bar = 20 μm. (**B**) Quantification of the colocalization of FS and PFF showed no significant difference (ns) compared to control using unpaired Student's t test. n = 6 for each condition (n = 12 total) collected from three independent experiments; mean ± SD. (**C**) U2OS cells were plated on coverslips and administered dextrans with different molecular weights (10,000; 70,000; 2,000,000) alongside PFF. Scale bar = 20 μm. (**D**) Quantification of dextran colocalization with PFF during uptake shows that their endocytic pathway is minimally overlapping, specifically in the case of 10,000 MW dextran; however, the internalization of dextran is much slower, especially with increasing molecular weight. n = 6 for each condition (n = 18 total) collected from three independent experiments; mean ± SD.

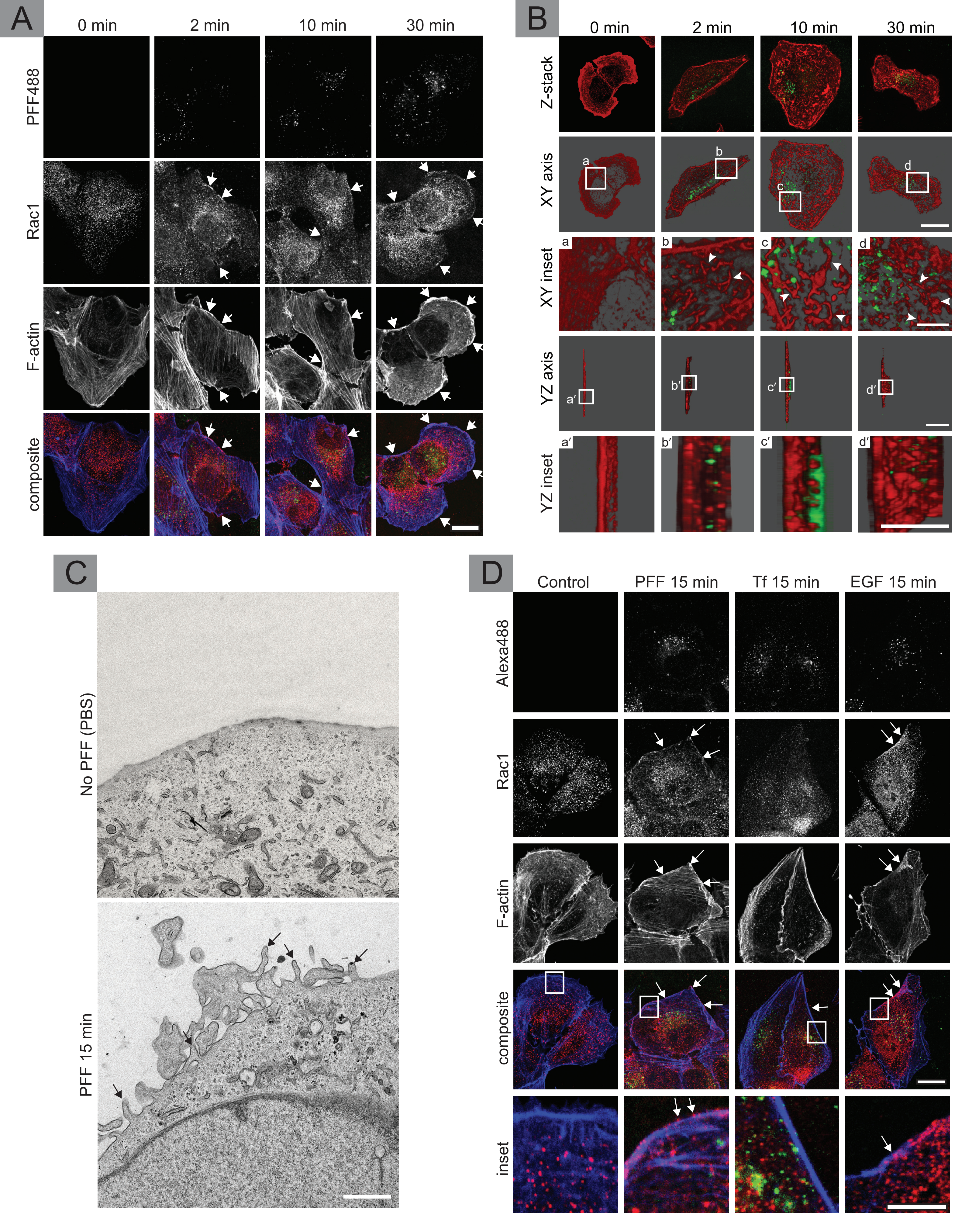


Bayati et al., Fig. S5

**Figure S5, related to Figure 4. Exposure to PFF leads to actin membrane ruffling and Rac1 recruitment.**

(**A**) Astrocytes were grown and mounted on coverslips. PFF (green) was added to each coverslip at 2 μg/ml and cells were incubated for 0, 2, 10, and 30 min at 37°C. Cells were washed with trypsin, fixed, permeabilized, and stained with Rac1(red) and F-actin (blue) antibodies. Arrows point to colocalization of F-actin and Rac1 at the cell surface, a sign of membrane ruffling. Scale bar = 20 μm. (**B**) Phalloidin (red) was used to stain F-actin in astrocytes with different PFF (green) incubation times. The Z-stacks were then combined to produce 3D images. Arrowheads point to actin ruffles. Scale bar = 20 μm and 5 μm for XY and YZ insets. (**C**) EM of astrocytes exposed to PBS or PFF for 5 min. Arrows point to membrane protrusions and ruffling. Scale bar = 1 μm. (**D**) Astrocytes were grown and mounted on coverslips. PFF, EGF, or Tf (all tagged with Alexa 488; green) were added to each coverslip at 2 μg/ml. Cells were then incubated for 0 or 15 min at 37°C. Cells were washed with trypsin, fixed, permeabilized, and stained with Rac1 (red) and F-actin (blue) antibodies. Arrowheads point to the colocalization of F-actin and Rac1 at the cell surface, a marker for membrane ruffling. Scale bar = 20 μm.

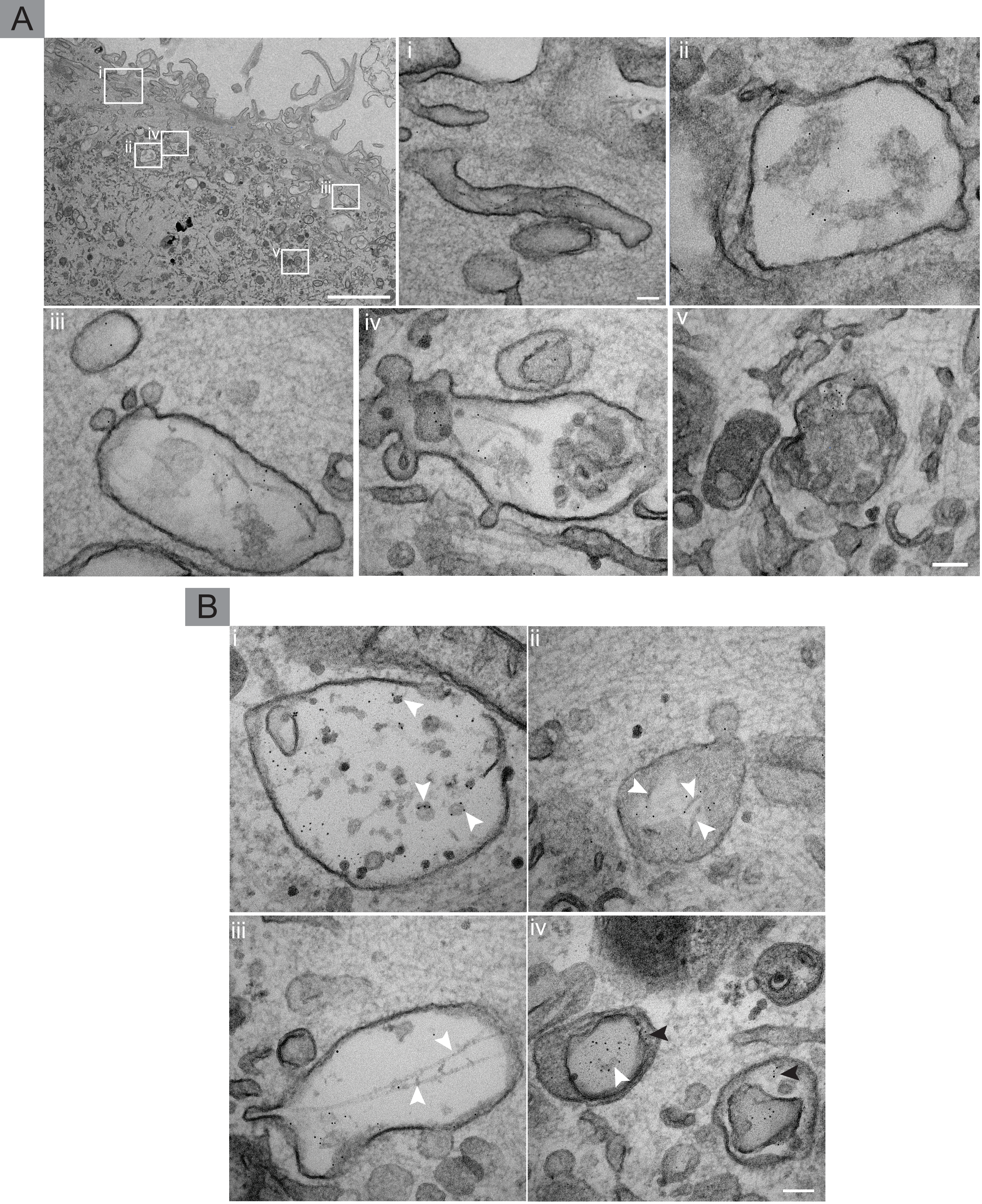


Bayati et al., Fig. S6

**Figure S6, related to Figure 6. PFF induces membrane ruffling at the cell membrane and is trafficked to multivesicular bodies via macropinosomes.**

(**A**) Astrocytes exposed to PFF-gold for 5 min showed PFF-gold localized to macropinosomes (**i-iii**). (**iv**) PFF is also found in multivesicular bodies. (**v**) PFF can also be found in electron dense multivesicular bodies and lysosomes. Scale bar = 2 μm for low magnification and 100 nm for all insets. (**B**) More examples of PFF located in various cellular compartments: astrocytes exposed to PFF-gold for 30 min. (**i**) Large multivesicular bodies, with many intraluminal vesicles with PFF on their surface (arrowhead), showing a medley of interactions between PFF and membranes. (**ii**) Visible PFF in the lumen of large vesicles (arrowhead). (**iii**) Long fibrils of α-syn (arrowhead) were found in macropinosomes at 30 min, suggesting that it takes longer for lengthy fibrils to enter the cell. (**iv**) PFF found both within (white arrowhead) and outside (black arrowhead) the lumen of large intraluminal vesicles. Scale bar = 100 nm.

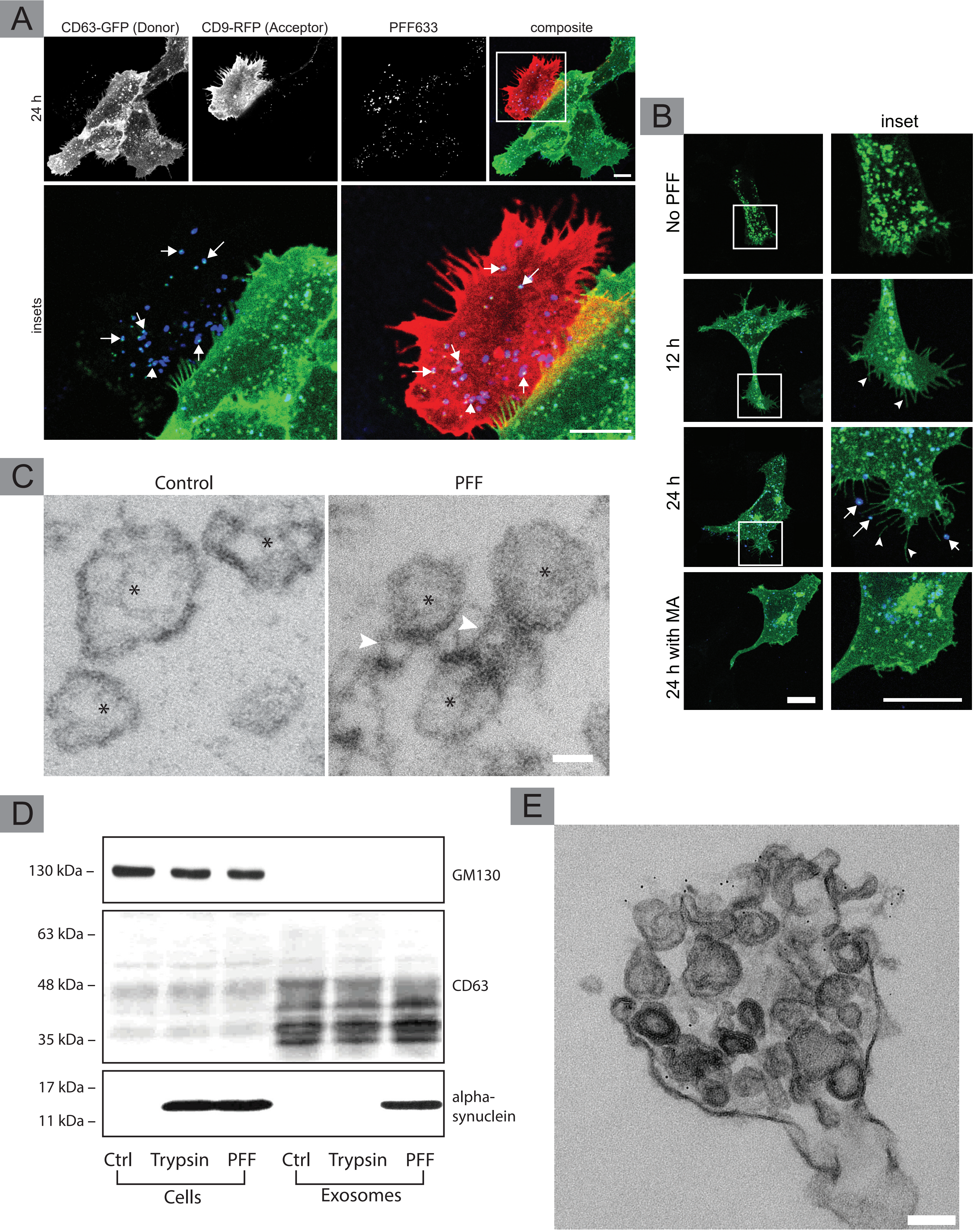


Bayati et al., Fig. S7

**Figure S7, related to Figure 7. CD63 positive exosomes play a role in the intercellular transmission of PFF from donor to acceptor cells.**

U2OS cells stably expressing CD63-GFP were exposed to PFF or PBS (vehicle control) for 24 h. CD63-GFP (donor cells) were then trypsin washed three times, pelleted, trypsinized again, pelleted, and PBS washed before being coplated with acceptor cells (PFF naïve cells stably expressing CD9-mCherry). Donor and Acceptor cells were then incubated for 24 h (**A**) 24 h samples show PFF (blue) and CD63 (green) fluorescence in CD9-positive (red) acceptor cells. The left insets show PFF and CD63 composite images without the CD9 channel, while the right insets show a closeup of the composite. Arrows point to compartments in acceptor cells that contain both CD63 and PFF. Scale bar = 20 μm for low magnification images and 10 μm for inset. (**B**) U2OS cells stably expressing CD63-GFP were exposed to PFF or PBS (vehicle control). Cells exposed to PFF (blue) showed the recruitment of CD63-GFP to membrane ruffles (arrowheads). At 24 h, cells readily showed exosomes (CD63 puncta not attached to the cell; arrows) colocalizing with PFF (blue) found near the end of CD63 ruffles. Cells incubated with PFF for 24 h but exposed to Manumycin (MA) showed very little CD63 ruffling. Scale bar = 20 μm for both low magnification images and insets. (**C**) U2OS cells exposed to PFF were trypsin washed three times and replated with regular media. 48 h following incubation, cell media was collected and underwent multiple centrifugation steps to isolate exosomes. A small portion of the exosome samples was then processed for EM. Exosomes (indicated by *) collected from cells exposed to PFF showed fibril-like structures on their surface. Scale bar = 20 nm. The remaining portion of the exosome samples were immunoblotted along with their cells of origin in **D**. The exosome samples were enriched in CD63 and were negative for GM130. Exosomes collected from PFF exposed cells were trypsinized and blotted along with the control and PFF exosomes. Like the control, trypsin exposed exosomes were not positive for α-syn. (**E**) Media from U2OS cells exposed to PFF-gold was collected and underwent centrifugation. The PFF gold sample pelleted at much lower centrifugation speeds, and the sample was processed for EM. Clusters of exosomes with PFF on their surface were found using EM. Scale bar = 100 nm.
